# Bulk and selective autophagy cooperate to remodel a fungal proteome in response to changing nutrient availability

**DOI:** 10.1101/2024.09.24.614842

**Authors:** Bertina Telusma, Jean-Claude Farre, Danica S. Cui, Suresh Subramani, Joseph H. Davis

## Abstract

Cells remodel their proteomes in response to changing environments by coordinating changes in protein synthesis and degradation. In yeast, such degradation involves both proteasomal and vacuolar activity, with a mixture of bulk and selective autophagy delivering many of the vacuolar substrates. Although these pathways are known to be generally important for such remodeling, their relative contributions have not been reported on a proteome-wide basis. To assess this, we developed a method to pulse-label the methylotrophic yeast *Komagataella phaffii* (*i*.*e. Pichia pastoris*) with isotopically labeled nutrients, which, when coupled to quantitative proteomics, allowed us to globally monitor protein degradation on a protein-by-protein basis following an environmental perturbation. Using genetic ablations, we found that a targeted combination of bulk and selective autophagy drove the vast majority of the observed proteome remodeling activity, with minimal non-autophagic contributions. Cytosolic proteins and protein complexes, including ribosomes, were degraded via Atg11-independent bulk autophagy, whereas proteins targeted to the peroxisome and mitochondria were primarily degraded in an Atg11-dependent manner. Notably, these degradative pathways were independently regulated by environmental cues. Taken together, our new approach greatly increases the range of known autophagic substrates and highlights the outsized impact of autophagy on proteome remodeling. Moreover, the resulting datasets, which we have packaged in an accessible online database, constitute a rich resource for identifying proteins and pathways involved in fungal proteome remodeling.

## INTRODUCTION

Organisms have evolved conserved strategies to cope with fluctuations in available nutrient sources, including altering their proteome through a combination of modulated protein synthesis, protein degradation, and cell growth-driven protein dilution (Advani and Ivanov 2019; Bonny et al. 2021; Zatulovskiy et al. 2022; Reddien 2024). When facing changes in nutrient availability, including deficiencies, these pathways work collectively to deplete cells of now-superfluous proteins, thereby allowing for the reallocation of resources towards the synthesis of proteins required for viability in the new environmental condition. In yeast, such protein degradation is largely driven by the proteasome, a compartmentalized AAA energy-dependent protease, and the vacuole – a cellular compartment defined by a single membrane and filled with largely nonspecific hydrolases capable of degrading and recycling proteins, lipids, and nucleic acids (Sala et al. 2017).

The ability to degrade proteins is a fundamental aspect of an organism’s adaptation to changing conditions and this degradation is carefully regulated at multiple levels to protect against the toxic effects of nonspecific protein turnover (Lu et al. 2017; Pohl and Dikic 2019). Indeed, proteasomal degradation requires that protein substrates be recognized and unfolded by a regulatory complex that typically requires that substrates bear a polyubiquitin modification (Dikic 2017). Since proteasomal substrates must be mechanically unfolded and translocated through a narrow axial channel in the regulatory particle, proteasomal degradation is thought to primarily process single polypeptides sequentially (Dikic 2017). In contrast, protein complexes, aggregates, condensates, and cellular organelles often rely on the process of macroautophagy, hereafter referred to as autophagy, for targeting to, and subsequent degradation in, the lysosome/vacuole (Pohl and Dikic 2019; Dikic 2017). Through autophagy, substrates are engulfed by a double-membraned vesicle that later fuses with the vacuole where encapsulated substrates are degraded. In some instances, this encapsulation occurs non-selectively in a process known as bulk autophagy that, in fungi, strictly depends on the transmembrane protein Atg9 (He et al. 2006, 2008; Mari et al. 2010). Additionally, cells can utilize receptor proteins to selectively tether specific cargos to the core autophagic machinery, thereby facilitating their degradation while minimally compromising the rest of the cellular content. In yeast, this “selective autophagy” process relies on Atg9, the autophagy scaffolding protein Atg11, and cargo-specific receptor proteins, including Atg30 and Atg32, which target peroxisomes and mitochondria for selective autophagic degradation, respectively (Farré et al. 2008; Farré and Subramani 2016; He et al. 2006; Kanki et al. 2009; Motley et al. 2012; Okamoto et al. 2009).

The methylotrophic yeast *Komagataella phaffii*, also known as *Pichia pastoris*, is particularly well-suited to studying proteomic responses to changing nutrients as it can be grown on a variety of carbon sources, including methanol, glucose, and oleate, each of which requires a uniquely tailored proteome (Gould et al. 1992; Hou et al. 2022; Moser et al. 2017; Wriessnegger et al. 2007). As such, transitioning between carbon sources necessitates altered levels of key metabolic pathways to efficiently utilize the available carbon compounds, with these alterations often achieved through proteome remodeling. For instance, when provided the reduced C_1_-compound methanol as the sole source of carbon and energy, *K. phaffii* grows robustly using a methanol utilization pathway that is distributed between the cytosol and a specialized membrane-bound organelle known as the peroxisome (Hartner and Glieder 2006; Rußmayer et al. 2015; Zhan et al. 2023). Indeed, growth of *K. phaffii* in methanol induces a massive proliferation of peroxisomes and associated methanol utilizing enzymes, including alcohol oxidase (AOX), which is targeted to the peroxisome (Chang et al. 1999; Sibirny 2016; Zhang et al. 2017). In contrast, upon adaptation to alternative carbon sources, such as glucose, these peroxisomes are rapidly degraded in the vacuole through a selective form of autophagy known as pexophagy (Tuttle et al. 1993; Tuttle and Dunn 1995), and the cell concomitantly upregulates glycolysis and other glucose-utilizing pathways (Nazarko et al. 2008).

Whereas the pathways responsible for degrading specific substrates in response to changing environmental conditions have been identified for a small subset of individual substrates in *K. phaffii*, we lack a clear, systems-wide understanding of the contributions of the ubiquitin-proteasome system (UPS) and of selective or bulk autophagy to proteome remodeling, particularly during times of nutrient adaptation or starvation. Here, we develop a general approach to quantify proteome dynamics by coupling stable isotope pulse-labeling to quantitative mass spectrometry in *K. phaffii*. We then employ this assay in the context of growth condition perturbations and mutations targeting key genes involved in the aforementioned degradation pathways. The resulting quantitative resource describes time-resolved dynamics of a combined ∼3,200 proteins across four growth perturbations in six different genetic backgrounds. Further, we detail exemplar uses of this resource to assess the relative contributions of bulk autophagy, selective autophagy, and non-autophagic processes to cellular adaptation in changing environmental conditions. We find that bulk autophagy drives degradation of most cellular compartments under conditions of nutrient stress, with selective autophagy supporting degradation when cells are grown in steady-state conditions in methanol. We further observe distinct selective autophagy pathways, with pexophagy occurring rapidly relative to mitophagy under our stress conditions. Importantly, we discover that these degradative pathways do not compete for a limited resource, as ablation of either pathway has no impact on the rate of the other.

## RESULTS

### Protein degradation drives proteome remodeling during glucose adaptation

*K. phaffii* grows robustly on a variety of primary carbon sources, including glucose, methanol, and oleate (Gould et al. 1992; Riley et al. 2016), and does so by adjusting its proteome composition in each growth media (Hou et al. 2022). Indeed, when comparing the proteome of cells grown in methanol to that of cells grown in glucose, we identified ∼270 proteins whose levels changed more than two-fold using quantitative mass spectrometry (**Supplementary Figure 1A,B**). Specifically, we observed large changes in levels of proteins linked to methanol and glucose metabolism (**Supplementary Figure 1C,D**), consistent with the concept of metabolic specialization (Pereira et al. 2018; Xiao et al. 2023). To understand how cells transitioned from a methanol-adapted proteome to one suited for growth on glucose, we assayed the response of methanol-specific proteins to glucose adaptation via pulse-labeling coupled to quantitative mass spectrometry (PL-qMS) (Chen et al. 2012; Jomaa et al. 2014; Davis et al. 2016), which we hypothesized would allow us to independently monitor protein synthesis and degradation on a per-protein basis across large swaths of the proteome.

Briefly, we grew wild-type *K. phaffii* strain GS115 in “light” ^12^C-labeled, ^14^N-labeled methanol media (M+N). At time 0, this M+N media was rapidly replaced with “heavy” ^13^C-labeled glucose media that lacked nitrogen (G-N) (**Figure 1A**; see Methods). Quantitative mass spectrometry of tryptic peptides isolated from these cells revealed that the stable isotope pulse distinguished newly synthesized proteins (^13^C-labeled, post-pulse) from those present pre-pulse (^12^C-labeled, pre-pulse) based on the isotope envelope in their mass spectra (**Figure 1B**). Notably, throughout the time course, a fixed volume of culture was harvested, which allowed for direct comparison of the pre-pulse abundance over time without the need to consider dilution by growth. We quantified ∼3,200 across three replicate time courses and observed that the abundance of many ^12^C-labeled peptides (pre-pulse) decreased as a function of time following the pulse, consistent with protein degradation (**Figure 1C**). Although most proteins underwent degradation upon this nutrient transition, a minority remained stable (**Supplementary Table 1**), including those annotated as cell-wall resident (Burgard et al. 2020). This observation aligned with the notion of cell wall inaccessibility to lysosomal and proteasomal degradation and indicated that the protein degradation we observed across the vast majority of the proteome did not result from cell lysis.

**Figure 1.**
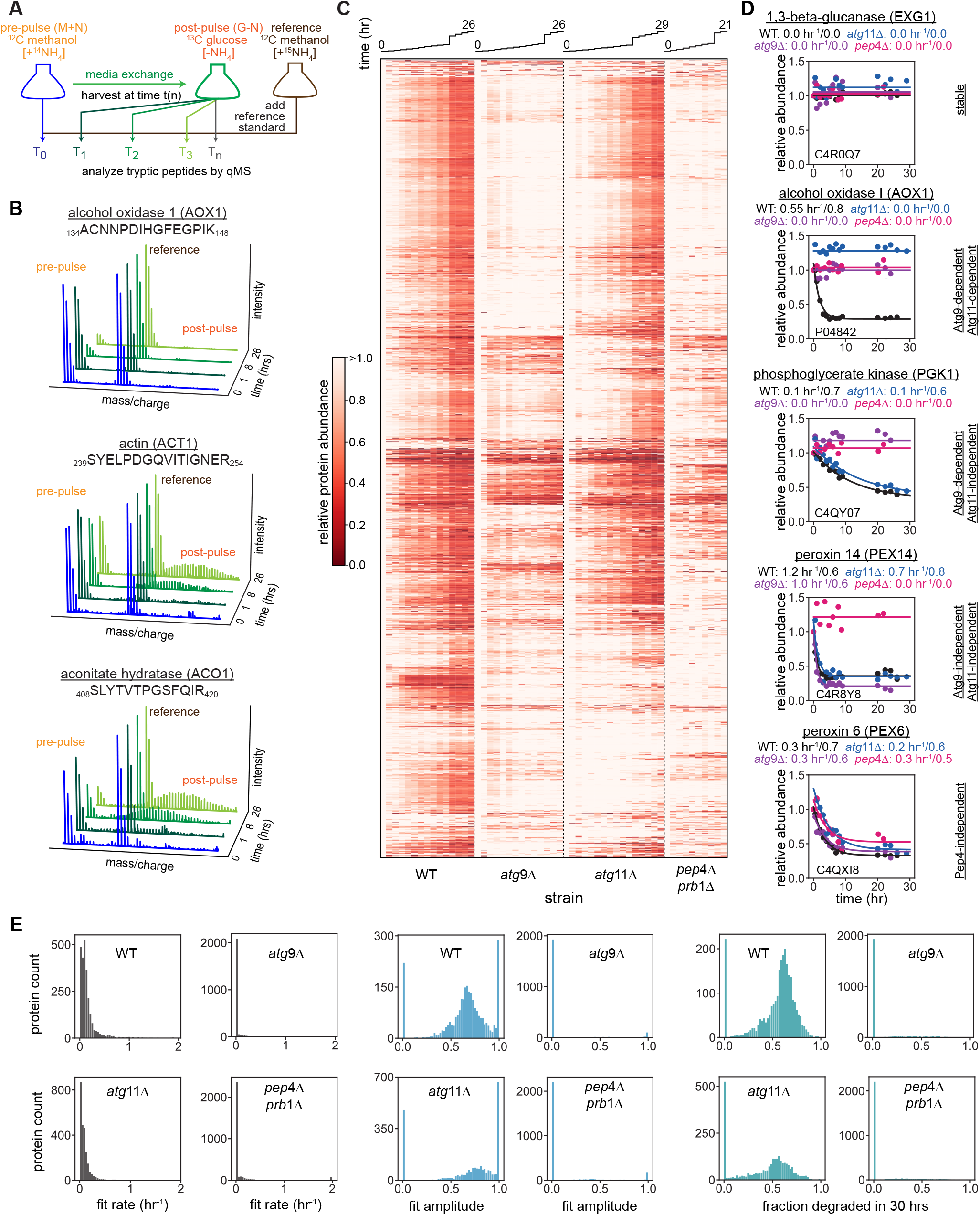
Stable isotope pulse-labeling coupled to mass spectrometry quantifies protein degradation. **(A)** Schematic depiction of pulse-labeling experimental design. Media composition pre- and post-pulse, including isotopically labeled species noted. At each timepoint, a fixed volume of culture was added to a fixed reference standard, which was ^15^N-labeled. **(B)** Exemplar mass spectra highlighting proteins degraded (top), synthesized (middle), and simultaneously degraded and synthesized (bottom). Pre-pulse, reference, and post-pulse isotope envelopes noted. A subset of timepoints (0, 2, 8, and 26 hours) from wild-type cells are plotted. (**C**) Clustered heatmap of proteome degradation compared between wild type, *atg*9Δ, *atg*11Δ, *pep*4Δ*prb*1Δ strains transitioning from growth in M+N to G-N media. Columns represent individual timepoints (see Methods) and rows represent individual proteins. For each peptide, relative abundance value at time n was calculated as the pre-pulse peptide abundance relative to that in the reference isotope channel, normalized to that at time 0. The abundance of each isotope species was estimated from the precursor (MS^1^) signal of the M0, M1, and M2 isotopomers using Spectronaut. That is: (pre-pulse_n_/reference)/(pre-pulse_0_/reference). Protein relative abundance is calculated as the median relative abundance across all uniquely mapped peptides. Timeponts plotted as a linechart above clustermap. (**D**) Representative fits highlighting the different modes (annotations at right of plots) of degradation captured in C. Kinetic traces are fit as: abundance(*t*) = A_deg_*exp(-*k*_*deg*_*t) + B (*eq. 1*, see Methods), with fit parameters *k*_*deg*_/A_deg_ noted above each plot. Uniprot protein ID and protein common names are listed. **(E)** Distributions of the rate (left) and amplitude (middle) parameters fit for each protein in each condition as in D. Using these fit parameters, a univariate measure of the fraction of pre-pulse material degraded in 30 hours for each protein was determined, and the distribution of this estimated parameter is plotted for each strain (right).

By carefully inspecting each isotopic envelope, we could additionally monitor protein synthesis, observing proteins that were: 1) exclusively synthesized; 2) simultaneously synthesized and degraded; and 3) degraded in the absence of new synthesis (**Figure 1B**, top, middle, bottom, respectively). We observed that proteins central to methanol utilization were robustly degraded with minimal synthesis, whereas proteins that facilitate glucose utilization were rapidly synthesized and degraded at rates that increased their levels on balance (**Supplementary Figure 2A-C**). Taken together, we found that the vast majority (∼90%) of the quantifiable proteome underwent degradation, and that many of the robustly synthesized proteins were related to glucose metabolism.

### Atg9-dependent bulk autophagy drives proteome remodeling

We next assessed the role of autophagy in the large-scale protein degradation we observed by repeating this nutrient shift/PL-qMS in autophagy-deficient *atg*9Δ cells. Most (2,266/2,549) proteins were stabilized in the Atg9 deletion background, consistent with Atg9-dependent autophagy driving much of the proteome remodeling observed in wild-type cells. Notably, when this experiment was repeated in cells lacking vacuolar proteases (*i*.*e. pep*4Δ*prb*1Δ), the results were similar to that in *atg*9Δ cells (**Figure 1C**). To determine the functional consequences of autophagy inhibition, we measured the growth rates of WT and *atg*9Δ cells in our various media. Observed growth rates were virtually indistinguishable between these strains in either methanol (M+N) or glucose (G+N) media, and they initially grew similarly when shifted from methanol (M+N) to glucose media lacking nitrogen (G-N). However, after ∼5 hours in G-N media, growth of *atg*9Δ cells ceased, whereas growth persisted in WT cells for at least an additional 10 hours, highlighting the functional impact of autophagy on nutrient adaptation. This effect was not glucose-dependent, as a similar dependence on Atg9 for cell growth was observed as cells transition from M+N to M-N media (**Supplementary Figure 3**).

### Rates of autophagic degradation vary broadly between protein substrates

To quantify degradation proteome-wide, we modeled the time-dependent change in protein abundance as a single-exponential decay process (see Methods), producing apparent first-order degradation rate constants (rate, *k*_deg_), and the fraction of initial material degraded (amplitude, A_deg_) for each protein. In wild-type cells, the half-lives of proteins varied widely, from ∼1 hour to over 130 hours (corresponding to *k*_deg_ ranging 0.8 hr^-1^ to 0.005 hr^-1^), and amplitudes ranged from 0.2 to 1. Interestingly, many proteins were only partially degraded, (*i*.*e*. A_deg_ < 1), consistent with proteolytic protection of these proteins via potential mechanisms such as differential subcellular localization (Bensimon et al. 2020; Simpson et al. 2022), post-translational modification (Botti-Millet et al. 2016; Lee et al. 2023; Wani et al. 2015), or exhaustion of degradative capacity (Ben-Zvi et al. 2009; Hipp et al. 2019; Kumar et al. 2023; Santra et al. 2019). In *atg*9Δ and *pep*4Δ*prb*1Δ cells, we observed sharp reductions in the observed degradation rates and amplitudes that collectively had profound impacts on the fraction of initial material degraded during a 30-hr timeframe (**Figure 1C-E**). We next defined Atg9-dependent substrates as those whose kinetic fit reported that more than 20% of the starting material was degraded after 30 hours in WT cells, and less than 20% was degraded in the same time span in the *atg*9Δ strain. Although most protein degradation was ablated in these autophagy-deficient cells, a small number of proteins were degraded in an Atg9-independent fashion (**Figure 1D**). Interestingly, these Atg9-independent substrates separated into two different groups: 1) those whose degradation relied on vacuolar proteases consistent with macroautophagy-independent vacuolar degradation (**Supplementary Table 2**), and 2) those degraded in a Pep4/Prb1-independent fashion, consistent with degradation by the ubiquitin-proteasome system or other non-vacuolar proteases (**Supplementary Table 3**). Interestingly, this group included the proteins PEX1, PEX5 and PEX6, which are thought to be part of the peroxisomal PEX5 receptor recycling/RADAR pathway (Ma et al. 2013), and PEX5 is a reported proteasomal substrate when this recycling pathway is blocked (Platta et al. 2007). Notably, strain-dependent substrate degradation profiles, and their associated genetic dependencies were largely consistent across replicates (**Supplementary Figure 4**).

### Atg11-dependent selective autophagy is active during proteome remodeling

To assess the relative contributions of bulk and selective autophagy, we next quantified growth rates and protein degradation in *atg*11Δ cells, as Atg11 is thought to be a critical scaffold supporting selective autophagy in yeast (He et al. 2008; Mari et al. 2010; Zientara-Rytter and Subramani 2020). This experiment revealed that: 1) wild-type and *atg*11Δ cells grew similarly in all media tested (**Supplementary Figure 3**); 2) that most protein substrates were degraded similarly in the two strains when cells transitioned from M+N to G-N media; and 3) that a cluster of proteins required both Atg9 and Atg11 for their degradation in response to this media change (**Figure 1C-E**). suggesting that selective autophagy drives their turnover. Indeed, when inspecting canonical bulk autophagy substrates (Wang et al. 2020; Welter et al. 2010), such as the cytosolic protein phosphoglycerate kinase (PGK1), we observed robust degradation in WT and *atg*11Δ cells but not in the *atg*9Δ strain, consistent with the Atg11-independent nature of bulk autophagy (**Figure 1D**). In contrast, peroxisomal substrates such as alcohol oxidase and catalase required both Atg9 and Atg11, consistent with pexophagy, a well characterized form of selective autophagy (Farré et al. 2008; Tuttle and Dunn 1995), driving their degradation. Notably, some peroxisomal turnover also occurs by micropexophagy, wherein the yeast vacuole directly engulfs peroxisomes without involving the double-membraned autophagosome (Oku and Sakai 2016). However, as this process is also Atg9-dependent (Chang et al. 2005), we were unable to distinguish the relative contributions of micro- and macropexophagy.

### Nutrient adaptation promotes distinct compartment-specific degradation patterns

To further characterize the selectivity of degradation in response to this nutrient shift, substrates were classified based on their cellular compartments using Gene Ontology annotations (Aleksander et al. 2023; Ashburner et al. 2000) (see Methods). We then compared strain-specific degradation profiles of proteins exclusively localized to the nucleus, cytoplasm, endoplasmic reticulum, Golgi apparatus, mitochondria, peroxisome, and plasma membrane. This analysis revealed that, with the exception of peroxisomes and mitochondria, most proteins in the other cellular compartments we analyzed were robustly degraded in an Atg9-dependent, Atg11-independent fashion, suggesting that bulk autophagy plays a significant role in the degradation of diversely localized proteins (**Figure 2A**). For example, proteins from the Golgi apparatus and those from the ER, which together form a central hub for the secretory pathway, were largely degraded in an Atg9-dependent, but Atg11-independent fashion, with similar kinetics observed for proteins throughout the secretory pathway (**Supplementary Figure 5A**). Likewise, cytosolic ribosomal proteins were largely degraded in an Atg9-dependent, Atg11-indepdendent fashion, consistent with bulk autophagic turnover. Notably, however, slow degradation of large subunit proteins was observed in both *atg*9Δ and *pep*4Δ*prb*1Δ cells, indicating contributions of non-vacuolar proteolysis to the turnover of the large subunit specifically (**Supplementary Figure 5B**). This observation stands in contrast with prior reports of a strict autophagic requirement for degradation of large ribosomal subunits in yeast (Kraft et al. 2008).

**Figure 2.**
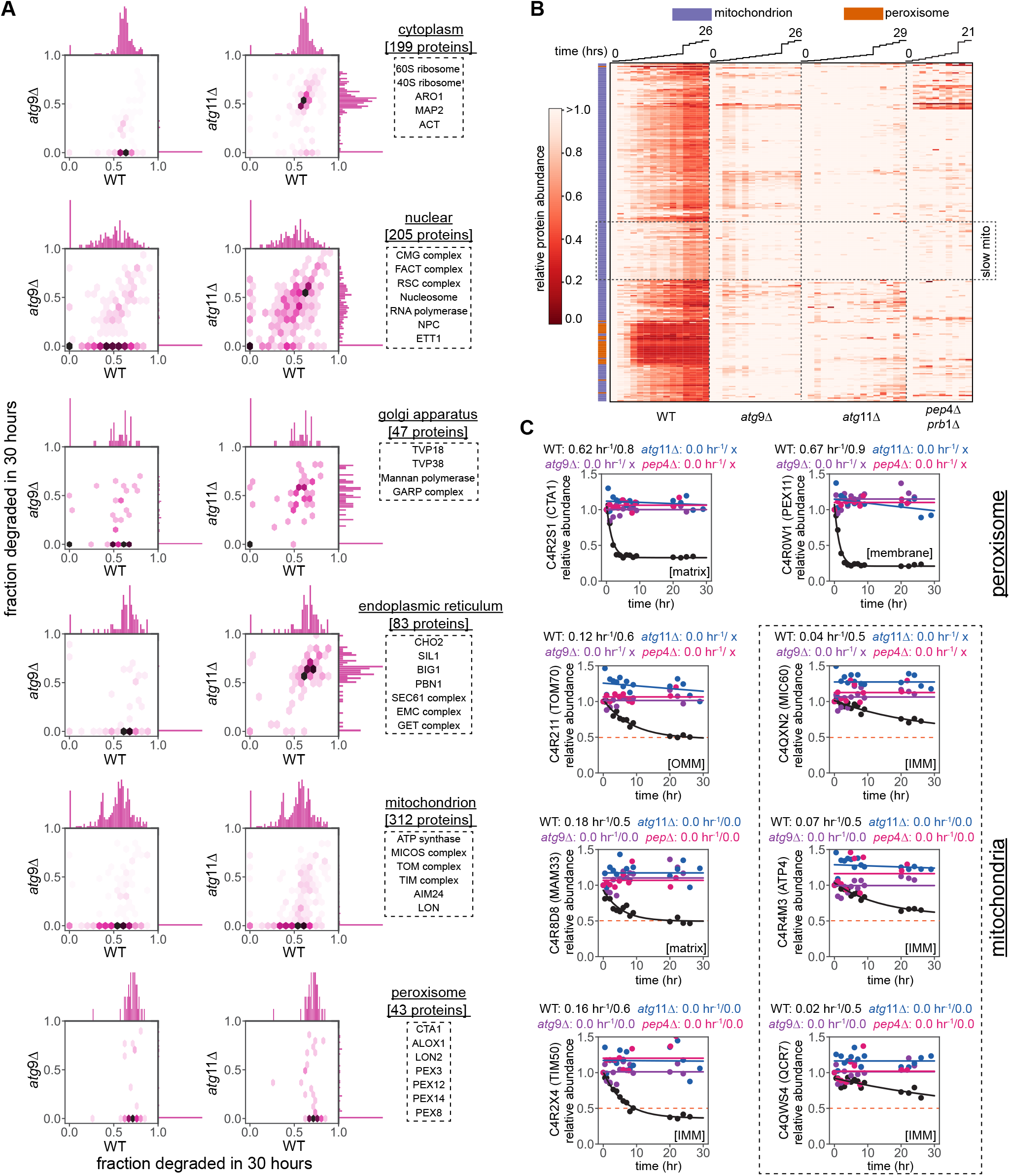
Selective autophagy drives degradation of some, but not all, organelle-localized proteins. **(A)** Proteins were grouped based on annotated subcellular localization (see Methods), and the fit kinetic model for each protein was used to calculate the fraction of initial material degraded in 30 hours in wild-type (WT), *atg*9Δ, and *atg*11Δ cells. This value is compared between genetic backgrounds as a density hexplot with marginal distributions shown on top and right axes. Exemplar protein or protein complexes from each compartment are noted in dashed boxes. **(B)** Clustered heatmap following Figure 1 depicting annotated mitochondrial (purple) and peroxisomal (orange) proteins subject to selective autophagic degradation. A cluster of mitochondrial proteins degraded more slowly than typical mitochondrial proteins are outlined. **(C)** Degradation profiles and fit kinetic models for representative proteins localized to the peroxisome (top) or mitochondria (bottom), with sub-organellular localization in brackets (OMM: outer mitochondrial membrane; IMM: inner mitochondrial membrane). Kinetic traces are fit following *eq. 1* (see Methods), with fit parameters *k*_*deg*_/A_deg_ noted. Uniprot IDs and protein common names are listed. The dashed box highlights a subset of slowly degraded IMM proteins.

Degradation in these compartments contrasted with that in the peroxisomes and mitochondria, including mitochondrial ribosomes, where Atg11-dependent selective autophagy was prevalent (**Figure 2B, Supplementary Figure 5C**). Notably, peroxisomal degradation was rapid relative to that of mitochondria. Within the mitochondria, we observed differential degradation rates for various protein complexes, with many members of the F_1_/Fo ATP synthase, cytochrome C oxidase, MICOS, NADH dehydrogenase, and pyruvate dehydrogenase complexes degraded more slowly than other mitochondrial proteins (**Figure 2C, Supplementary Figure 6, Supplementary Table 4**). We additionally observed Atg9- and Prb1/Pep4-independent degradation of mitochondrial respiratory chain proteins COX13 and COQ4 (**Supplementary Figure 7A, Supplementary Table 5**), which were likely targets of AAA proteases present in the mitochondria (Arlt et al. 1998; Korbel et al. 2004).

We also observed diverse patterns of genetic dependencies for degradation of proteins involved in peroxisome biogenesis. Specifically, proteins strictly localized to the peroxisomal membrane, such as PEX2, PEX3, PEX11, and PEX12, were degraded in a selective autophagy-dependent manner, likely captured through pexophagy. In contrast, many cytosolic proteins implicated in peroxisome biogenesis, including PEX5, which was previously suggested to be a selective autophagy substrate (Wang et al. 2020), were degraded through Atg11-independent pathways, with some also exhibiting Atg9- and Pep4/Prb1-independent degradation (**Supplementary Figure 7B, Supplementary Table 5**).

### Bulk and selective autophagy collaborate to degrade entire metabolic pathways

In *K. phaffii*, proteins required for methanol utilization are distributed in the cytosol and peroxisome (van der Klei et al. 2006; Yurimoto et al. 2005; Hartner and Glieder 2006; Sakai et al. 1998), and we speculated that this difference in cellular localization might necessitate different pathways to clear those substrates. Indeed, while steady-state measurements showed that both methanol-enriched cytoplasmic and peroxisomal proteins are expressed at similar levels (**Supplementary Figure 1C**), our nutrient switch/PL-qMS revealed they are degraded through different pathways. For instance, peroxisomally localized enzymes were rapidly removed through selective autophagy whereas those in the cytoplasm were slowly removed through bulk autophagy (**Supplementary Figure 8, Supplementary Table 5**). Triose-phosphate isomerase (TPI) was the sole example of a protein canonically thought to localize to the cytosol that was degraded via selective autophagy, raising the possibility of this enzyme bearing a cryptic peroxisome targeting sequence (Freitag et al. 2012) in *K. phaffii*. Interestingly, no methanol utilization-related proteins were degraded without Atg9, highlighting the primacy of autophagy relative to proteasomal degradation in this context.

### Basal autophagy is active under steady-state conditions

The extensive proteome remodeling observed in response to changing nutrients prompted us to investigate whether autophagy also actively degraded proteins under steady-state growth conditions. To assess this, cells were grown under steady-state conditions in M+N media, and a PL-qMS assay was performed (see Methods). Under these steady-state conditions, we observed minimal protein degradation, with the vast majority of detected proteins exhibiting proteolytic stability up to 50 hours (**Figure 3A, Supplementary Figure 9A**). These stable proteins included markers of bulk autophagy (*e*.*g*. phosphoglycerate kinase PGK1), consistent with a lack of bulk autophagy in this growth media (**Figure 3A**). Nevertheless, peroxisomal turnover was detected as evidenced by the selective autophagic degradation of peroxisomal proteins such as peroxiredoxin (PMP20), catalase (CTA1), and the peroxin PEX11, whereas methanol-enriched cytosolic proteins were stable (**Figure 3, Supplementary Figure 9B**). Similarly, mitochondrial proteins were largely stable in these steady-state conditions (**Supplementary Figure 9C**).

**Figure 3.**
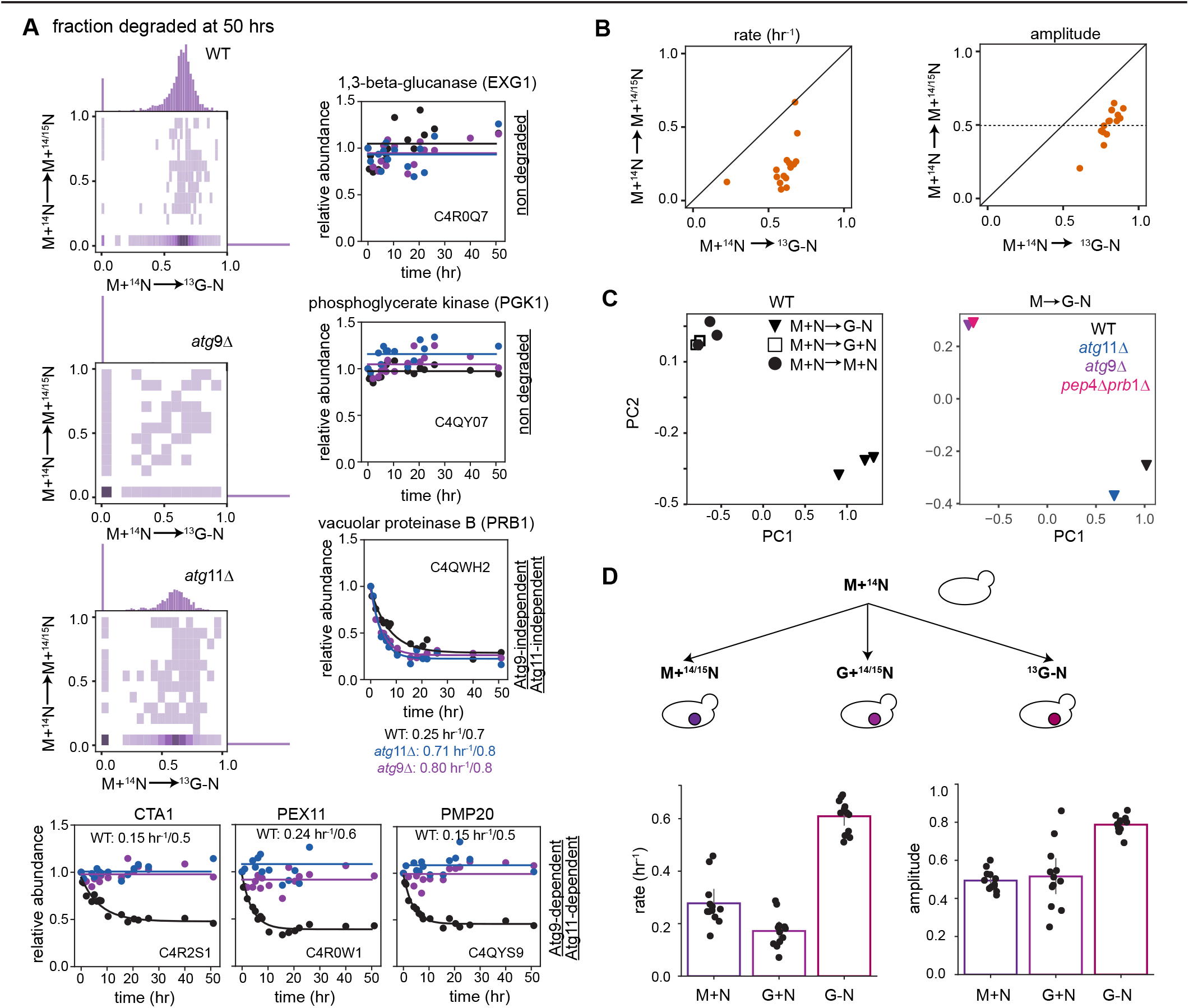
Peroxisomes are degraded in steady-state growth conditions. **(A)** Degradation across the proteome compared between cells grown at steady-state in nitrogen-rich methanol media (M+^14^N →M+^14/15^N) to those transitioning to glucose media without nitrogen (M+^14^N→^13^G-N). In each condition, the fit kinetic model for each protein was used to calculate the fraction of initial material degraded in 30 hours in wild-type, *atg*9Δ, and *atg*11Δ cells. This value is compared between different media conditions post-pulse as a density hexplot with marginal distributions shown on top and right axes. Degradation profiles measured in steady-state growth in methanol are shown for exemplar substrates exhibiting noted genetic dependencies for degradation. Kinetic traces are fit as: abundance(*t*) = A_deg_*exp(-*k*_*deg*_*t) + B (*eq. 1*), with fit parameters *k*_*deg*_/A_deg_ noted. Uniprot protein ID and protein common names are listed. **(B)** The rate (left) and amplitude (right) of 15 peroxisome-localized proteins compared between cells transitioning from methanol media to glucose media without nitrogen (M+^14^N→^13^G-N) and those grown in steady state conditions in methanol media (M+^14^N→M+^14/15^N). **(C)** Principal component analysis (PCA) of the fit degradation rates and amplitudes for each protein measured in wild-type cells undergoing the media transitions: M+N to M+N (circles), M+N to G+N (squares), or M+N to G-N (triangles), with each mark noting an independent biological replicate (left). PCA as described above, but now used to globally compare degradation in WT, *atg*9Δ, *atg*11Δ, and *pep*4Δ*prb*1Δ cells transitioning from growth on M+N to G-N media (right). **(D)** Schematic of media transitions (top) and fit degradation rates and amplitudes of the peroxisomal proteins, compared across the different media transition conditions (bottom), with dots denoting individual peroxisomal proteins and boxes marking average values.

Furthermore, pexophagy occurred at a slower rate and to a lesser extent in steady state conditions compared to that observed under nutrient-shift induced conditions (**Figure 3B**). Surprisingly, under these steady-state conditions fit amplitudes were ∼0.5 for these peroxisomal proteins (*i*.*e*. ∼50% of these peroxisomal proteins would never be degraded), suggesting an uncharacterized mechanism by which cells can protect subsets of peroxisomal proteins from the basal levels of selective autophagy we observed.

### Nitrogen starvation is the primary driver for the observed proteome remodeling

Next, we assessed the isolated contributions of changing the carbon source and of removing nitrogen to the proteome remodeling we observed as cells transitioned from M+N to G-N media. We first measured protein turnover as cells transitioned from M+N to G+N media (see Methods), finding that much of the proteome was stable, with the overall pattern of degradation as analyzed by principal components most similar to cells grown in steady state M+N conditions. The minimal degradation observed was similar to that seen in *atg*9Δ and *pep*4Δ*prb*1Δ cells transitioning between M+N to G-N media (**Figure 3C**). Moreover, like in steady-state growth conditions, selective turnover of the peroxisomes was observed, and the rate and extent of degradation was reduced relative to that seen as cells adapted to G-N media (**Figure 3D**).

To directly test whether the nitrogen starvation signal was the primary driver of our previously observed massive proteome remodeling, we assessed turnover as cells transitioned from M+N to M-N media. As previously observed, when we limited nitrogen and changed the carbon source, large differential proteome remodeling was induced under nitrogen starvation stress alone, with bulk autophagy contributing substantially to this degradation (**Figure 4A**). As in the M+N to G-N transition, we found examples of stable proteins, those degraded via bulk autophagy, selective autophagy, or independent of autophagy (**Figure 4B**). Selective autophagic turnover of peroxisomes and mitochondria was again observed, with peroxisomes degraded more rapidly than mitochondria (**Supplementary Figure 10A**). Additionally, degradation of proteins involved in peroxisome biogenesis, methanol utilization, and the secretory pathway exhibited varied genetic dependencies (**Supplementary Figure 10B-C**).

**Figure 4.**
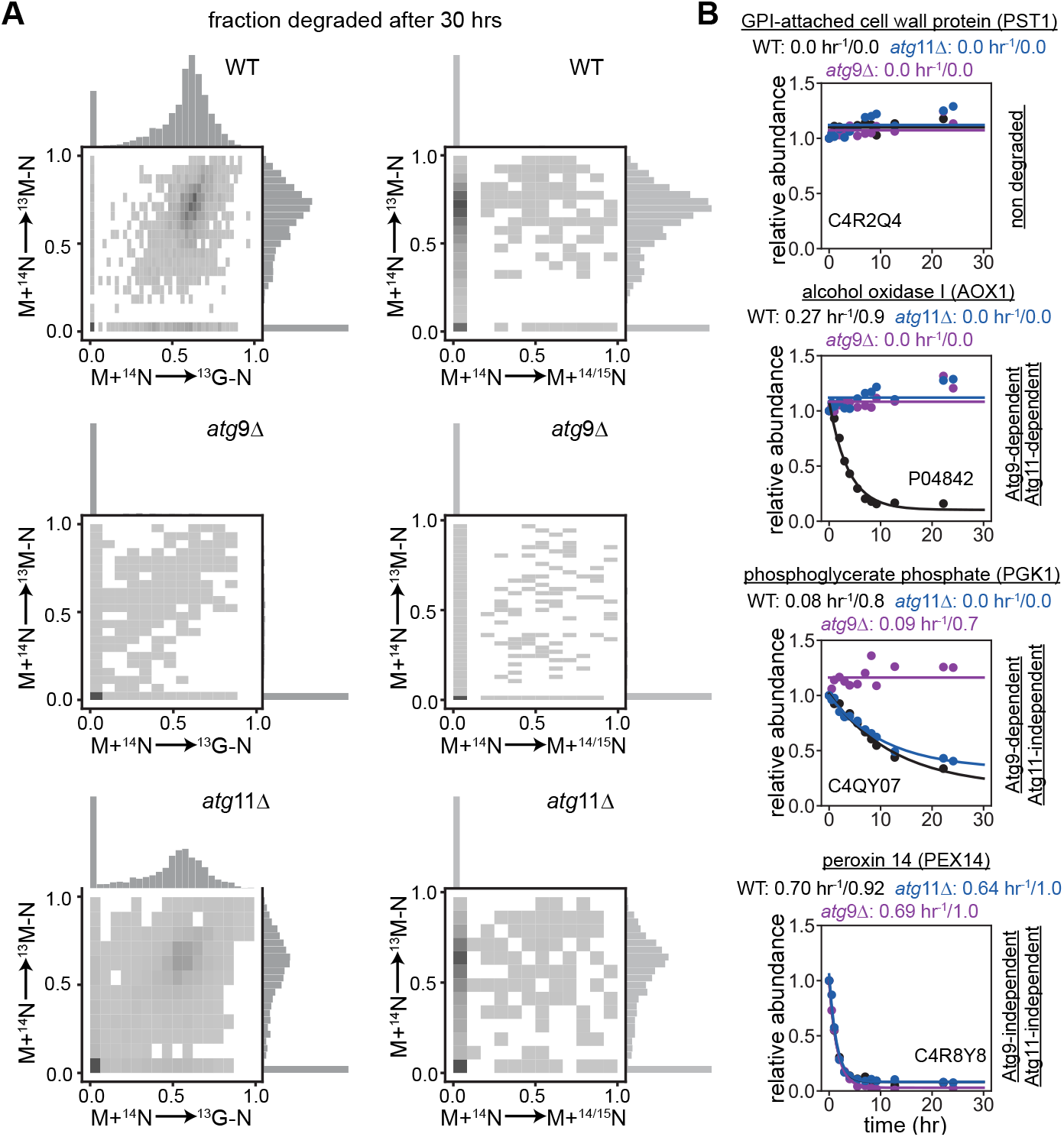
Nitrogen starvation is sufficient to promote proteome-wide turnover. **(A)** Proteome-wide degradation compared between cells transitioning from M+^14^N to ^13^G-N media and cells transitioning from M+^14^N to ^13^M-N media. In each condition, the fit kinetic model for each protein was used to calculate the fraction of initial material degraded in 30 hours in wild-type (top), *atg*9Δ (middle), and *atg*11Δ (bottom) cells. This value is compared between different media conditions post-pulse as a density hexplot with marginal distributions shown on top and right axes. **(B)** Different degradation profiles exhibiting noted genetic dependencies for degradation as cells transitioned from M+^14^N to ^13^M-N media. Kinetic traces are fit as: abundance(*t*) = A_deg_*exp(-*k*_*deg*_*t) + B (*eq. 1*, see Methods), with fit parameters *k*_*deg*_ and A_deg_ noted above each plot. Uniprot protein ID and protein common names are listed.

### Rates of mitophagy and pexophagy are independently governed

The disparate degradation rates we observed suggested that the mitochondrial and peroxisomal degradation pathways could be competing for a single rate-limiting resource – Atg11-associated autophagy initiation complexes, for example – or, instead, could be independently regulated and prioritized (**Figure 5A**). To explore this idea, we asked whether inhibiting degradation of either mitochondria or peroxisomes could stimulate the degradation of the other, as would be predicted by the limiting resource model. In *K. phaffii*, the receptor protein Atg30 marks peroxisomes for autophagic degradation (Farré et al., 2008), whereas receptor protein Atg32 targets mitochondria to autophagosomes (Kanki et al. 2009; Okamoto et al. 2009). Performing our PL-qMS assay in *atg*30Δ cells resulted in a near-complete loss of pexophagy, with minimal impacts on bulk autophagy or mitophagy. Similarly, loss of Atg32 impaired mitophagy but did not affect the turnover of peroxisomes (**Figure 5B-C, Supplementary Figure 10A**), indicating that a shared resource is not limiting degradative flux of either pathway. Instead, these findings suggest that fluxes through the pexophagy and mitophagy pathways are independently regulated, potentially at the level of activation of the selective autophagy receptors themselves (Zientara-Rytter et al. 2018; Tanaka et al. 2014; Kanki et al. 2013).

**Figure 5.**
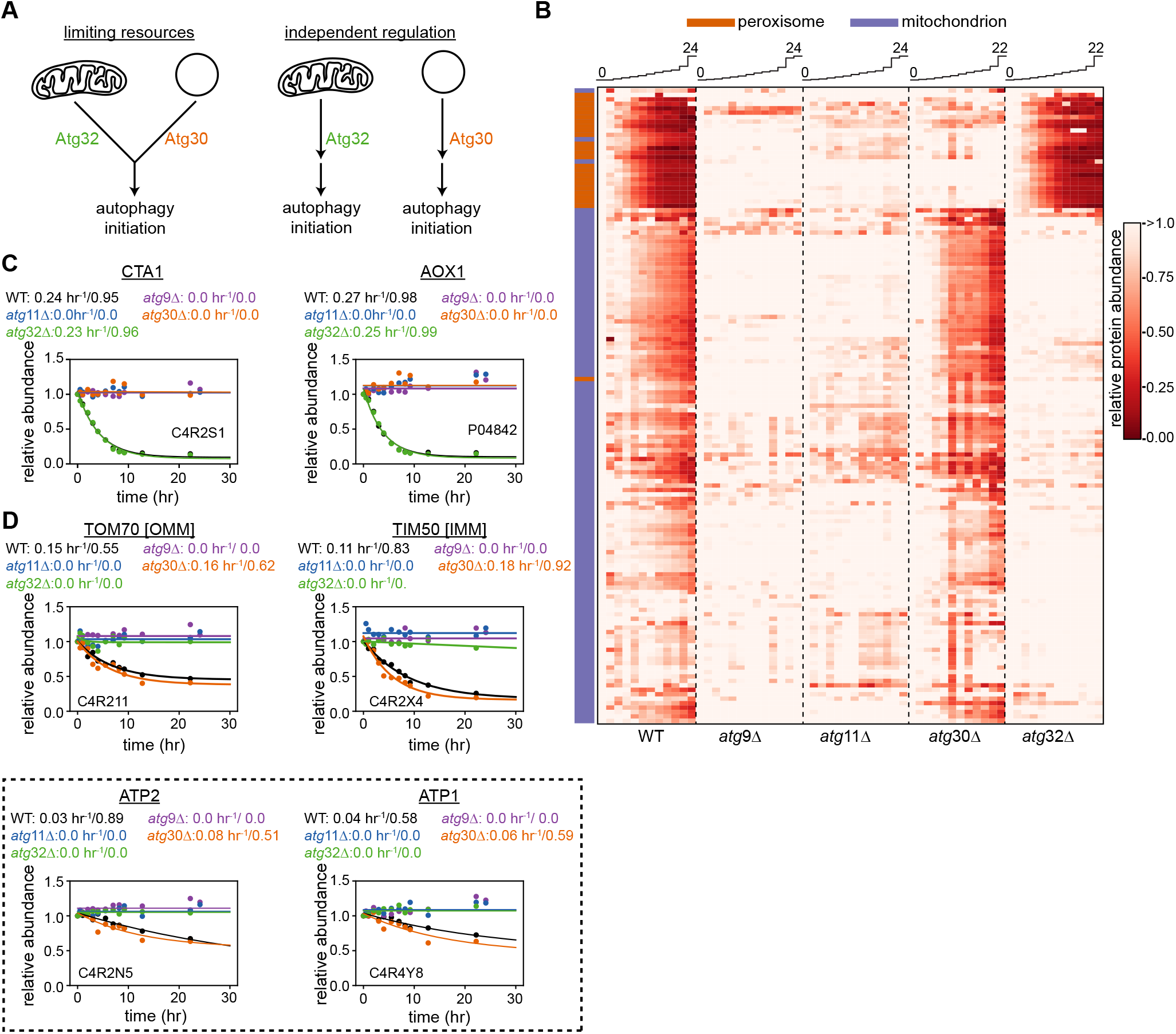
Selective autophagic pathways are independent regulated. **(A)** Two hypothetical models for how peroxisomes and mitochondrial are degraded through autophagy. In the shared limiting resource model (left), the receptors Atg30 and Atg32 compete for the same limiting autophagic machinery or resource such that the presence of one pathway is expected to slow the other. In the independent regulatory model (right) the pathways are not in competition and are not predicted to impact one another. **(B)** Clustered heatmap plotting the relative abundance of peroxisomal (orange) or mitochondrial (purple) proteins over time in wild-type, *atg*9Δ, *atg*11Δ, *atg*30Δ, and *atg*32Δ cells transitioning from M+N to M-N media. **(C-D)** Degradation profiles and fit kinetic models (*eq. 1*, see Methods) of representative peroxisomal (C) or mitochondrial (D) proteins, with Atp synthase related proteins, which are degraded relatively slowly, outlined. Uniprot IDs and protein common names are listed, with fit parameters *k*_*deg*_ and A_deg_ noted above each plot.

## DISCUSSION

It has long been appreciated that nutrient limitations profoundly influence protein degradation pathways (Reddien 2024), including the ubiquitin proteosome system and bulk and selective autophagy. However, in most instances, our understanding of the relative contributions of these different pathways to overall proteolytic flux has been limited, in part due to limitations of the techniques employed. For example, western blotting (Kraft et al. 2008; Meiling-Wesse et al. 2002; Welter et al. 2010) and fluorescence assays (Ichimura et al. 2000; Kaizuka et al. 2016; Kirisako et al. 2000; Koepke et al. 2020) only provide targeted measurements of a handful of substrates. They additionally rely on the specificity of affinity reagents or the ability to genetically integrate tags without impacting target function to accurately quantify physiological protein levels. Mass spectrometry enables near proteome-wide tag-free measurements (Hickey et al. 2023) but, as typically implemented, can only measure steady-state protein levels, which obscures proteins whose synthesis is balanced with degradation. For example, we observed significant peroxisomal degradation (and balanced synthesis) under steady state (M+N) growth conditions, which would have been overlooked with traditional western, fluorescence, or mass spectrometry-based approaches.

Alternative approaches to identify proteins targeted by different proteolytic pathways include proximity assays such as IP-MS (Behrends et al. 2010) or APEX-based proximity labeling (Zellner et al. 2021), with each producing a catalog of potential substrates. These proximity-based methods in isolation cannot, however, distinguish between co-localization that leads to degradation and that which does not, and these methods fail to provide turnover rates, degradation kinetics, or the fraction of a given substrate that is degraded under specific conditions, each of which can be critical for assessing proteome remodeling.

Within the autophagy field, overall autophagic activity levels are often assayed using reporters, such as the ratio of lipidated to non-lipidated Atg8-family proteins, proteolytic cleavage of a marker protein fused to an autophagic substrate, or quantitation of autophagosomes via transmission electron microscopy or fluorescent puncta associated with autophagosome formation (Klionsky et al. 2021). Notably, these assays cannot assess the specific substrates targeted by autophagy and, in the absence of additional modulations aimed at revealing “autophagic flux”, it can be challenging to distinguish an increase in formation of autophagosomes from a failure of autophagosome/lysosome fusion (Chang et al. 2017; Loos et al. 2014). Moreover, none of these assays provide a kinetic rate for autophagy generally, let alone on a per-substrate basis.

Our PL/qMS approach provides a global view of protein degradation, allowing one to measure both degradation rates and the total fraction of proteins degraded on a per-substrate basis. As such, it can be used as a quantitative, functional readout of which proteins are degraded in response to an environmental change, but it cannot directly ascribe the proteolytic pathway utilized. Thus, we have coupled our approach to genetic ablation of key genes required for various forms of autophagy. In *K. phaffii*, this method was effective, as the degradation of many proteins exhibited switch-like responses to these manipulations, with protein degradation completely ablated in the cognate knockout strain. This switch-like behavior contrasted with that observed in cultured human cells where Cui and colleagues observed many protein substrates that could flexibly access either lysosomal or proteasomal degradation pathways (Cui et al. 2024).

In designing our method, we were concerned that our isotope labeling scheme, which requires three mass-distinguishable labeling patterns to independently monitor protein synthesis and degradation would limit protein coverage. However, when coupled with advances in instrumentation (Kelstrup et al. 2018) and data analysis (Chi et al. 2018), we successfully detected and quantified ∼3,200 proteins of the *K. phaffii* proteome, which consists of ∼5,000 encoded proteins (The UniProt Consortium 2023), only a subset of which are expected to be expressed in any particular condition. This extensive coverage facilitated detailed tracking of subcellular compartments and their dynamics, allowing for deep comparisons between the various nutrient adaptation and genetic backgrounds we compared. Moreover, the resulting measurements constitute a rich resource to interrogate the regulation and dynamics of protein turnover in different conditions. These data are available through an interactive web application (jhdavislab.org/datasets) where users can inspect the degradation profiles of these proteins and assess how these profiles change in *atg*9Δ or *atg*11Δ cells.

To interrogate the pathways supporting adaptive response to nutrient stress, we subjected *K. phaffii* to a stringent stress condition by changing the media carbon and simultaneously depriving cells of nitrogen. Our aim was to induce stress responses, which we expected would lead to dynamic alterations in protein degradation and gene expression, as well as nutrient scavenging and cellular metabolism. Our findings emphasize that when *K. phaffii* adapts to nitrogen deprivation, either with or without a concomitant change in the carbon source, it initiates massive remodeling of the proteome, with a large but incomplete fraction of the proteome targeted for degradation. The remodeling is predominately driven by autophagy, as evidenced by the substantial stabilization of the proteome in *atg*9Δ cells and limited Atg11-dependence. Notably, ablation of bulk autophagy resulted in a severe growth defect, presumably as a result of the cell’s inability to recycle nutrients in the absence of Atg9-dependent bulk autophagy.

Bulk autophagic degradation targeted both cytosolic substrates and those associated with the secretory pathway, as evidenced by the similar degradative rates and genetic dependencies of proteins in the lumen or membrane of the ER, Golgi, and vesicles transiting between these compartments. Nuclear-localized proteins exhibited a more complex degradation dependence, with different proteins displaying genetic dependencies consistent with bulk autophagy, selective autophagy, or non-autophagic degradation, highlighting the complex interplay between the UPS system and autophagy in this compartment (Li and Nakatogawa 2022; Enam et al. 2018).

In conditions of acute nutrient stress, *K. phaffii* employs both bulk and selective autophagy to restructure its proteome. Our study specifically highlights the simultaneous activation of two selective autophagy pathways, mitophagy and pexophagy, with pexophagy degrading substrates rapidly relative to mitophagy. Additionally, we noted variations in the degradation kinetics of mitochondrial proteins with a key subset of proteins, including F_O_/F_1_ ATP synthase subunits, degraded more slowly than other mitochondrially localized proteins degraded via mitophagy. The simple topological organization of the mitochondria did not explain the observed degradation patterns, as we observed similar rates of degradation for proteins localized to the outer membrane or to the matrix. This lack of a topological dependence suggested that some protein complexes in the mitochondria may be more stable to vacuolar proteolysis, or otherwise less susceptible to autophagic degradation. In contrast to that in the mitochondria, selective autophagic degradation of peroxisomal proteins was more uniform, with both membrane-associated and matrix-localized proteins undergoing relatively rapid selective autophagic degradation.

A recent study in cultured human cells proposed that in some conditions, selective autophagy pathways are in competition and, specifically, that upregulated pexophagy limits aggrephagy and parkin-dependent mitophagy (Germain et al. 2024). The authors further posited that these pathways competed for limiting quantities of active ULK1, as pharmacological stimulation of ULK1 rescued the pexophagy-induced limitation in mitophagy. Interestingly, in our assays, selective autophagy pathways in yeast undergoing metabolic adaptation do not appear to be in competition for a limiting pool of initiation factors, as ablation of either pexophagy or mitophagy had no impact on the degradative capacity of the remaining pathway.

Whereas the bulk autophagic degradation we observed largely depended on nitrogen limitation, pexophagy was observed even under steady-state growth in methanol-containing media, where peroxisomes are continuously synthesized. Why might the cell simultaneously synthesize and degrade these proteins? We hypothesize that the observed turnover acts to maintain a healthy pool of peroxisomes, as the peroxisome-resident proteins are known to undergo oxidative damage as a byproduct of methanol metabolism, which produces significant levels of hydrogen peroxide (van der Klei et al. 2006; Aksam et al. 2009). However, if stochastic metabolism-associated damage were to trigger degradation of these proteins via pexophagy, one would expect simple first order kinetics, with all of the pre-pulse material eventually degraded. Instead, we observed that ∼50% of the pre-pulse peroxisome-associated proteins are stable, implying that they are either protected from damage, or otherwise sequestered from the degradative machinery. The mechanistic underpinnings of this observation, including whether the protected pool represents all peroxisomes within a subset of cells or a subset of peroxisomes within all cells, remain to be explored.

Taken together, this work provides a rich, quantitative resource to better understand how *K. phaffii* balances synthesis and degradation during steady state growth, and how the proteome is remodeled in response to nutrient changes, all with single-protein resolution. Moreover, it provides the requisite experimental tools and analytic approaches to quantitatively assess the roles of selective and bulk autophagy in supporting cellular adaptation to changing environments, including exposures to toxins, pathogens, and genotoxic, thermal, or oxidative stress.

## MATERIALS AND METHODS

### Yeast strains, media, and buffers

The GS115 *K. phaffii* (*P. pastoris*) strains used are listed in **Supplementary Table 1** and were kindly provided by the Subramani lab. All strains were cultured at 30°C in 125 mL baffled flasks at 225 RPM. Growth media (typically 5-25 mL) were defined as follows:

^**14**^**N Glucose Fed Medium (G+N):** 0.67% [w/v] yeast nitrogen base (YNB) without amino acids or ammonium sulfate (Research Products International, Y20040); 2.0% [w/v] glucose; 0.5% [w/v] ammonium sulfate (Fisher Scientific, BP212-212).

^**15**^**N Glucose Fed Medium (G+**^**15**^**N):** G+N with 0.5% [w/v] ^15^N-labeled ammonium sulfate (Cambridge Isotope Labs, NLM-713-25) replacing ^14^N ammonium sulfate.

^**13**^**C Glucose Nitrogen Starvation Medium (**^**13**^**G-N):** 0.67% YNB; 2.0% ^13^C glucose (Cambridge Isotope Labs, CLM-1396-10).

^**14**^**N Methanol Fed Medium (M+**^**14**^**N):** 0.67% YNB; 0.5% [v/v] methanol (Fisher Chemical, A412-4); 0.5% ammonium sulfate.

^**15**^**N Methanol Fed Medium (M+**^**15**^**N):** 0.67% YNB; 0.5% methanol; 0.5% ^15^N-ammonium sulfate.

^**13**^**C Methanol Nitrogen Starvation Medium (**^**13**^**M-N):** 0.67% YNB; 0.5% ^13^C methanol (Cambridge Isotope Labs CLM-359-5).

^**15**^**N Oleate Fed Medium (O+**^**15**^**N):** 0.67% YNB; 0.5% ^15^N-ammonium sulfate; 0.2% [v/v] oleic acid (Thermo Fisher); 0.02% [v/v] Tween-40 (Chem-Impex).

### Cell culturing and stable isotope pulse-labeling

#### Steady-state protein level measurements

For steady-state protein levels comparison between methanol and glucose, cells were separately grown in both ^14^N- and ^15^N-labeled G+N, M+N, and O+N media, and 5 mL of each culture were harvested at OD_600_ ∼ 0.5 via centrifugation 21,000 x g for 5 minutes. Following removal of supernatant, cells pellets were then washed with 500 µL of buffer A [20 mM Tris HCl (pH 7.5), 100 mM NH_4_Cl, 10 mM MgCl_2_, 0.5 mM EDTA, 6 mM 2-mercaptoethanol] and pelleted as before prior to storage at -80°C. During preparation of cells pellets, each ^14^N-labeled sample was resuspended in 2 mL of buffer A. A reference standard was generated by resuspending pellets bearing ^15^N-labeled cells in 2 mL of buffer A and mixing equal volumes each resuspension (G+^15^N; M+^15^N; O+^15^N). 50 µL of this mixture was then added to 50 µL of each of the ^14^N-labeled pellets, and these samples were then prepped for mass spectrometry as described below.

#### Pulse-labeling time courses

For steady-state pulse-labeling experiments in the presence of nitrogen, cells were initially grown at 30°C in pre-pulse media to an OD_600_ of ∼1.75, at which point they were pulse-labeled by the addition of an equal volume of pulse-media, which had been pre-warmed to 30°C. For starvation studies in the absence of nitrogen, nitrogen-containing pre-pulse grown cells were first pelleted at 3,000 x g for 5 min, washed with pre-warmed YNB buffer, pelleted at 3,000 x g for 5 min, and resuspended in pre-warmed starvation media. At each timepoint, 1 mL of culture was collected, harvested via centrifugation (21,000 x g, 5 min) at 4°C, washed with cold buffer A, and pelleted via centrifugation as above. Cell pellets were then rapidly frozen at -80°C prior to further analysis. In parallel, cells were grown in a ^15^N-labeled variant of the pre-pulse media, harvested at an OD_600_ of ∼1, and prepared and frozen as above.

Note that when monitoring proteome remodeling as cells transitioned from M+N to G+N media, cells were grown in “light” methanol media with ^14^NH_4_ and moved them to “heavy” G+N media bearing a 1:1 mixture of ^14^NH_4_:^15^NH_4_. This isotope pulse metabolically scrambles the nitrogen, resulting in partially labeled peptides that could be readily distinguished from the “light” precursors and from our reference standard grown in 100% ^15^NH_4_.

### Mass spectrometry sample preparation

After mixing collected samples with their cognate reference sample as described above, the mixture was immediately lysed by addition of trichloroacetic acid (TCA) to a final concentration of 13% (v/v). Samples were quickly frozen using liquid nitrogen and stored at -80°C for at least 1 hour or overnight if processed the following day. Sample preparation largely followed Sun et al. beginning with thawing samples and precipitating proteins by incubation in 13% TCA on ice for at least 2 hours (Sun et al. 2023). Samples were then centrifuged for 30 min at 4°C at 13,000 x g, and the supernatant discarded. The TCA precipitates were then washed with cold (4°C) acetone (500 µL) twice and dried at room temperature. Reduction of protein disulfide bonds was performed in 100 mM ammonium bicarbonate with 10 mM DTT at 65°C for 10 min, and protein alkylation was performed with 20 mM iodoacetamide at 30°C for 30 min in the dark. Proteins were then digested with 0.2 μg of trypsin/lysC (Promega) overnight at 37°C. Digested peptides were desalted using Pierce C18 columns or Thermo Fisher SOLA SPE plates following manufacturers’ protocols, dried down in a speedvac, and either stored at -80°C or resuspended in 20 µL sample buffer (4% acetonitrile, 0.1 formic acid) prior to analysis by LC-MS/MS.

### LC-MS/MS

Following desalting, tryptic peptides were separated using an Ultimate 3000 UHPLC system (Thermo Scientific) consisting of a trap column (C18, 75 µm ID x 2 cm, particle size 3 µm pore size 100 Å; Thermo Fisher Scientific, 164946) and an analytical column (75 µm ID x 50 cm, particle size 2 µm pore size 100 Å; Thermo Fisher Scientific, ES903). Peptides (2 µL, ∼1.5 µg) were loaded onto the trap column at a flow of 5 µL/min in a total volume of ∼20 uL. Peptides were washed on the trap column in loading buffer and then resolved on the analytical column using a 130 min linear gradient of 4%-30% buffer B (ACN, 0.1% FA) at a flow rate of 300 nL/min. Peptides were ionized by nano spray electrospray ionization and analyzed using an Thermo Scientific Orbitrap Exploris 480 (all M+N to M-N time courses; M+N to G-N *pep*4/*prb*1Δ time course) or HF-X (all other time courses) mass spectrometer as described below.

For data-dependent acquisition (DDA) runs, MS^1^ scans ranged from 350-1400 *m/z* and were collected at a mass resolution of 120,000 with AGC target and maximum injection time (maxIT) set at 3 × 10^6^ and 50 ms, respectively. The top 20 most abundant precursor ions were selected for subsequent MS^2^ scans and fragmented using 25% normalized collision energy (NCE). The fragment analysis (MS2) was performed at a resolution of 15,000, with an isolation mass window size of 2.2 m/z, AGC target and maxIT set at 1 × 10^5^ and 100 ms, respectively.

Data independent acquisition (DIA) MS^1^ scans followed those above as described for the DDA acquisitions. On the HF-X, MS^2^ scans were acquired using 25 windows spanning a mass range of 400-1250 *m/z* (**Supplementary Table 7**); precursors were fragmented using 28% NCE, and product ions were analyzed at a mass resolution of 30,000, with AGC and maxIT set at 1 × 10^6^ and 70 ms. On the Orbitrap Exploris 480, MS^2^ scans were collected using 10 m/z wide precursor windows overlapped by 2 m/z overlaps spanning the range 390-1390 m/z, with fragments generated using 28% NCE and analyzed at a resolution of 30,000 over a fixed 200-2,000 m/z range. Normalized AGC and maxIT were set at 2,000% and 70 ms, respectively.

### DDA data analysis

Raw DDA files were initially converted to the mzML format using MsConvert and subsequently searched within the trans-proteomic pipeline (Deutsch et al. 2010) using the Comet algorithm (Eng et al., 2013) against *Komagataella phaffii* Uniprot database for taxon ID 644223. The search parameters include 15 ppm precursor mass tolerance, 0.02 Da fragment mass tolerance, oxidized methionine as a variable modification, and carbamidomethyl cysteine as a static modification. The resulting peptide identifications were validated and scored using PeptideProphet and iProphet (Shteynberg et al. 2011). The resulting PepXML files were used to create a non-redundant spectral library in Skyline (MacLean et al. 2010). Raw DDA MS files were imported and searched against the spectral library for extraction of MS^1^ chromatographic peak areas. Skyline was used to qualitatively inspect peak shape, peak area, extracted chromatograms were inspected to ensure co-elution of light and heavy peaks.

#### DIA data analysis

Raw DIA files were processed using Spectronaut software (version 15) using DirectDIA mode (Bruderer et al. 2016). In brief, a spectral library was initially generated directly from DIA file by Spectronaut Pulsar, which incorporated a DDA library built from our DDA data. The settings for Pulsar and library generation were the same for both DIA and DDA files and were as follows: specific enzyme set to Trypsin/P, LysC, peptide length ranged 7 to 52; max missed cleavages=0; N-terminal M enabled; labeling included ^15^N at all nitrogens (2 channels utilized); fixed modification of carbamidomethyl Cys; variable modifications of Met oxidation and N-terminal acetylation; FDRs set to 0.01 and calculated at the PSM, peptide, and protein level; minimum fragment relative intensity 1%; 3-6 fragments kept for each precursor. For DIA analysis, Spectronaut search parameters were set as follows: mutation with NN predicted fragments to generate decoy, machine learning performed per run, precursor PEP cutoff=0.2; precursor q-value cutoff=0.01; single hit definition by stripped sequence. Using the report output, we exported MS^1^ intensities separately for channel1 (unlabeled) and channel2 (^15^N-labeled) for each precursor. The report file included: PG.Genes, PG.ProteinNames, EG.PrecursorId, PEP.MS1Channel1 and PEP.MS1Channel2.

### Steady-state data quantification

For steady-state protein level comparison of glucose-grown and methanol-grown *K. phaffii*, three biological replicates were processed as described above. DIA data was acquired and analyzed as previously described in Spectronaut. ^14^N (channel1) and ^15^N (channel2) MS^1^ intensities were separately exported for each detected peptide.

Using the exported MS^1^ intensities, ^14^N/^15^N ratios were generated for each identified precursor and the relative protein abundance was reported as the median relative peptide abundance across all peptides assigned to each protein. Additionally, for each growth condition, quantifiable proteins were normalized to their respective actin levels. Normalized log_2_ protein abundances were compared using a Student’s t-test, and the resulting p-values were corrected using the Benjamini-Hochberg multiple hypothesis correction procedure. A volcano plot was generated in Python, incorporating the adjusted p-values and log_2_-fold changes, with proteins exhibiting more than 2-fold changes with corrected p-values < 0.05 considered significant. Proteins involved in glucose metabolism and methanol metabolism were extracted from the statistically significant group and their relative abundances were compared.

### PL-qMS data quantification

For each peptide, its relative abundance at a timepoint was determined by normalizing the MS^1^ pre-pulse peptide (unlabeled) peak to the MS^1^ spike (labeled) peak. Relative protein abundances were defined as the median abundance across all peptides assigned to each protein. To quantify degradation on a protein-by-protein basis, individual protein abundance at different timepoints were fit to a single exponential function:

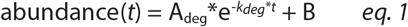

where, A_deg_, *k*_*deg*_, and B represent the amplitude, rate constant for degradation, and an offset, respectively. Note that in nearly all fits, A_deg_+B = 1, and with the exception of the most stable proteins, the data is well fit when B is fixed at 1-A. The fit quality was assessed using the sum of squared residuals, with poorly fit proteins manually inspected and discarded if the data was noisy. We next calculated the fraction of initial material degraded at t=30hrs (θ_30_) and defined proteins undergoing degradation as those for which at least 20% of their initial material was degraded (θ_30_>0.2) in 30 hours. Any protein with less than 20% fraction degraded (θ_30_<0.2) was defined as “stable”. This analysis was performed for the following backgrounds: WT, *atg*9Δ, *atg*11Δ, *atg*30Δ, *atg*32Δ, *pep*4Δ*prb*1Δ.

We then compared the extent of degradation across WT, *atg*9Δ, *atg*11Δ, *pep*4Δ*prb*1Δ strains to define different modes of degradation as follows:

<u>Atg9, Pep4-dependent</u>: WT θ_30_>0.2, *atg*9Δ θ_30_<0.2, *atg*11Δ *11Δ* θ_30_>0.2 *pep*4Δ*prb*1Δ θ_30_<0.2.

<u>Atg9, Atg11, Pep4-dependent</u>: WT θ_30_>0.2, *atg*9Δ θ_30_<0.2, *atg*11Δ θ_30_<0.2 *pep*4Δ*prb*1Δ θ_30_<0.2.

<u>Atg9, Atg11, Pep4-independent</u>: WT θ_30_>0.2, *atg*9Δ θ_30_>0.2, *atg*11Δ θ_30_>0.1, *pep*4Δ*prb*1Δ θ_30_>0.1.

<u>Atg9, Atg11-independent, Pep4-dependent</u>: WT θ_30_>0.2, *atg*9Δ θ_30_>0.2, *atg*11Δ θ_30_>0.2 *pep*4Δ*prb*1Δ θ_30_<0.2.

### Gene ontology analysis of quantified proteins

We used gene ontology (GO) annotations at the cellular localization category to assign subcellular localization data to proteins we analyzed, focusing on the terms nucleus, cytoplasm, endoplasmic reticulum, mitochondria, Golgi apparatus, peroxisome, vacuole, plasma membrane and cell wall. To characterize the mode of degradation of a specific compartment, we restricted our analysis to the subset of proteins that were strictly assigned to a single GO compartment. For instance, most quantified Golgi and the ER proteins were annotated to both compartments, but only those bearing a single annotation were analyzed when computing the organelle-scale mode of degradation. We categorized each compartment’s mode of degradation by inspecting the annotated proteins and assigned the mode of degradation as: 1) “non-degraded” if θ_30_ < 0.2 across WT, *atg*9Δ, *atg*11Δ, *pep*4Δ*prb*1Δ for the majority of the annotated proteins; 2) “bulk autophagy” if the majority of the annotated proteins were Atg9, Atg11, Pep4-dependent in their degradation; 3) “selective autophagy” if the majority of the annotated proteins were Atg9, Atg11, Pep4-dependent in their degradation. Note that even in compartments designated as autophagy targets, we observed non-autophagic degradation of a subset of proteins.

## DATA AVAILABILITY

The raw mass spectrometry data and summary tables will be publicly deposited upon publication. An interactive data browser is available at jhdavislab.org/datasets.

## AUTHOR CONTRIBUTIONS

B.T. and J.H.D. designed all experiments. B.T. performed all experiments and analyzed all data. J.C.F provided strains and guidance in culturing procedures and genetic manipulations. B.T. and J.H.D. wrote manuscript and generated figures. D.S.C. generated the interactive data viewer. J.H.D. supervised project. J.H.D. and S.S secured funding and edited the manuscript.

## ACKNOWLEDGEMENTS

We thank Laurel Kinman, April Lee, and Jen Kosmatka for helpful discussion and feedback. This work was supported by NIH grants R00-AG050749 (JHD), R01-GM144542 (JHD), DK41737 (SS), and NSF grants CAREER-2046778 (JHD), and GRFP (SMW).

## SUPPLEMENTARY FIGURES

**Supplementary Figure 1.**
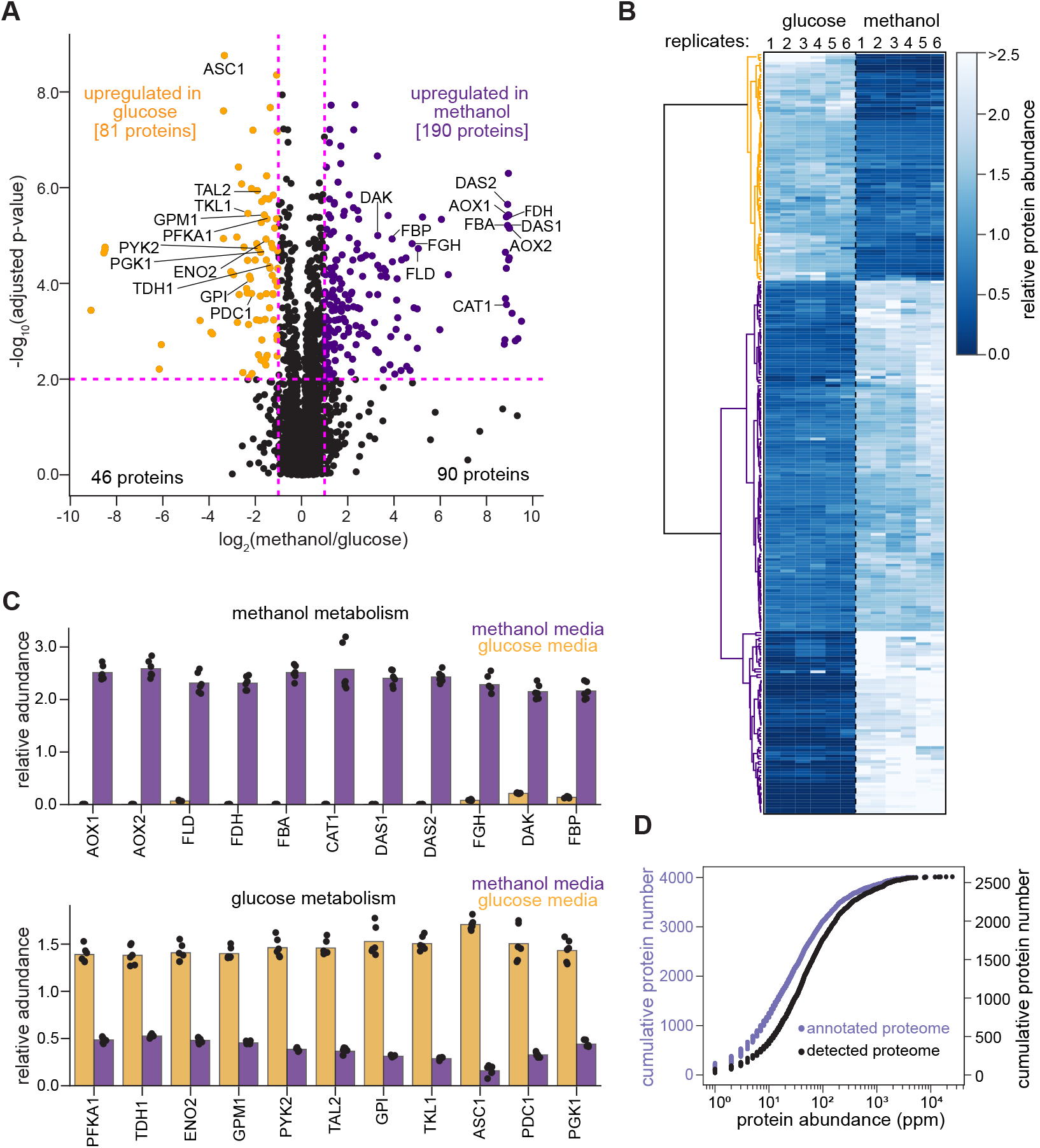
Carbon-source specific proteomes support growth of *K. phaffii*. **(A)** Volcano plot highlighting changes in protein levels between *K. phaffii* grown in methanol and glucose. Mean fold-change (log_2_ transformed) in protein abundance plotted against statistical significance (two-tailed t-test) calculated across six replicates for each condition (see Methods). Yellow and purple points indicate proteins of interest that exhibit a greater than 2-fold change in abundance (x-axis) with high statistical significance (y-axis; p-value ≤ 0.01). **(B)** Clustered heatmap depicting the relative protein abundance of the 271 proteins differentially expressed in glucose or methanol media. Clusters identifying glucose-enriched and methanol-enriched proteins colored in yellow and purple, respectively. **(C)** A subset of differentially expressed proteins involved in methanol (top) or glucose (bottom) metabolism. Each point represents an independent replicate (6 in total). **(D)** Cumulative frequency of protein abundance for proteins detected in our mass spectrometry assay (black) vs. that annotated in the *K. phaffii* proteome (purple). Protein abundance, which is reported as parts per million (ppm) was calculated for each *K. phaffii* protein with an ortholog in *S. cerevisiae* for which a ppm measurement was available PaxDb (Huang et al. 2023). Note that low abundance proteins are underrepresented in our assay.

**Supplementary Figure 2.**
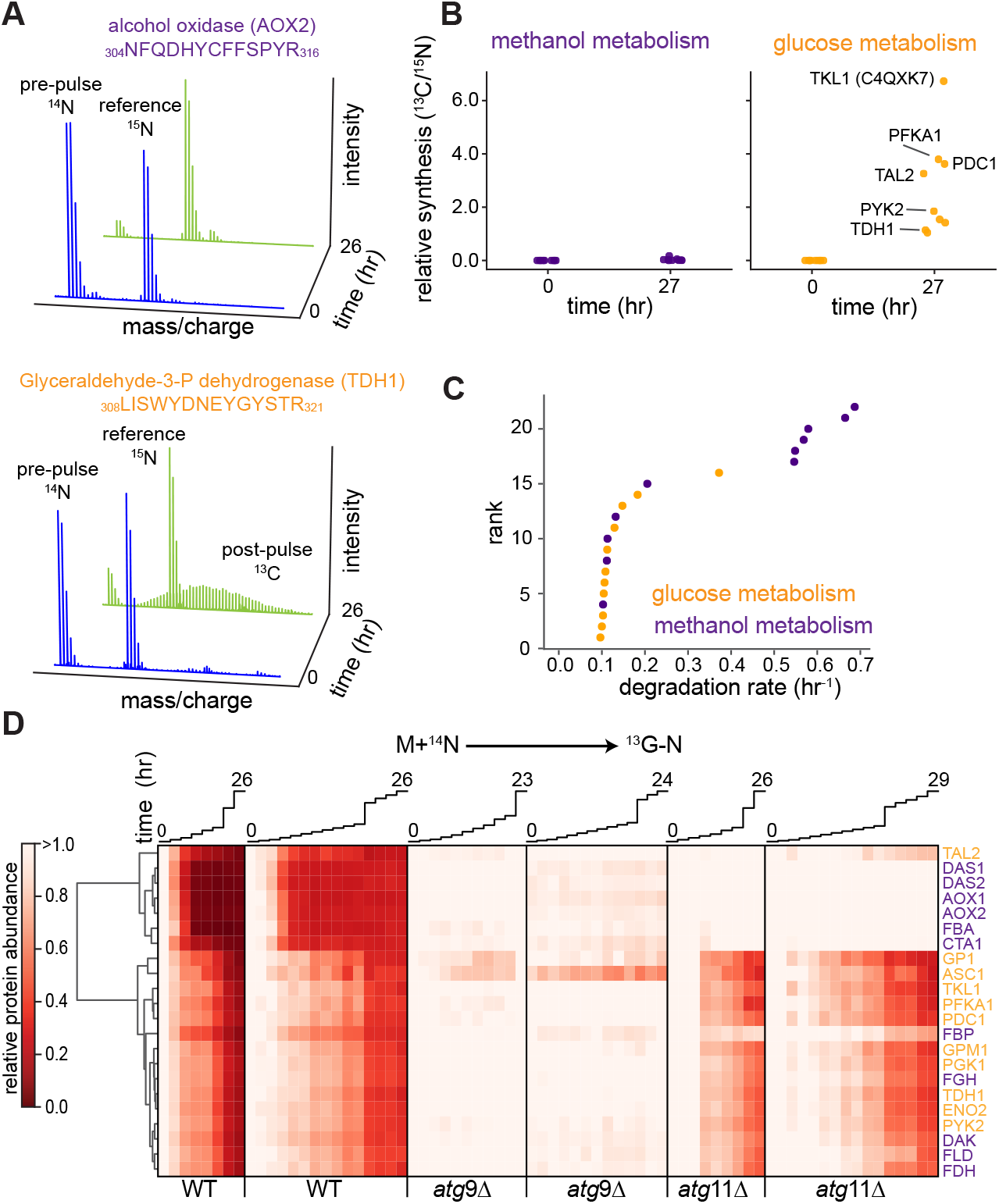
Differential synthesis and degradation of proteins involved in glucose and methanol metabolism. **(A)** Spectral representation of an exemplar peptide from alcohol oxidase (methanol enriched – AOX2; top) and glyceraldyhyde-3-phosphate dehydrogenase (glucose enriched – TDH1; bottom) at t=0 (blue) and t=26 hours (green) after shifting cells from growth in methanol (^12^C+^14^N) media to that in glucose media lacking nitrogen (^13^C-N). Note that post-pulse ^13^C-labeled peptides were visible in the TDH1 but not in the AOX2 spectra, and that the reference standard bearing a ^12^C+^15^N-labeled peptide was present in both. **(B)** Relative synthesis of proteins involved in methanol (purple) and glucose (yellow) metabolism (see Supplementary Figure 1C) following the media shift from (A). Relative synthesis quantified using the ratio of ^13^C- to ^15^N-labeled peptides at timepoints 0 and 27 hours. **(C)** Cumulative distribution of degradation rates measured for proteins involved in methanol (purple) and glucose (yellow) metabolism following media shift from (A). **(D)** Clustered heatmap depicting degradation profile of proteins from (C) across two replicates in WT, *atg*9Δ and *atg*11Δ cells. Relative protein abundance at each timepoint calculated as ratio (^12^C+^14^N; pre-pulse)- to (^12^C+^15^N; reference)-labeled peptides, normalized to that calculated at 0 hours. Columns represent individual timepoints, which are plotted above.

**Supplementary Figure 3.**
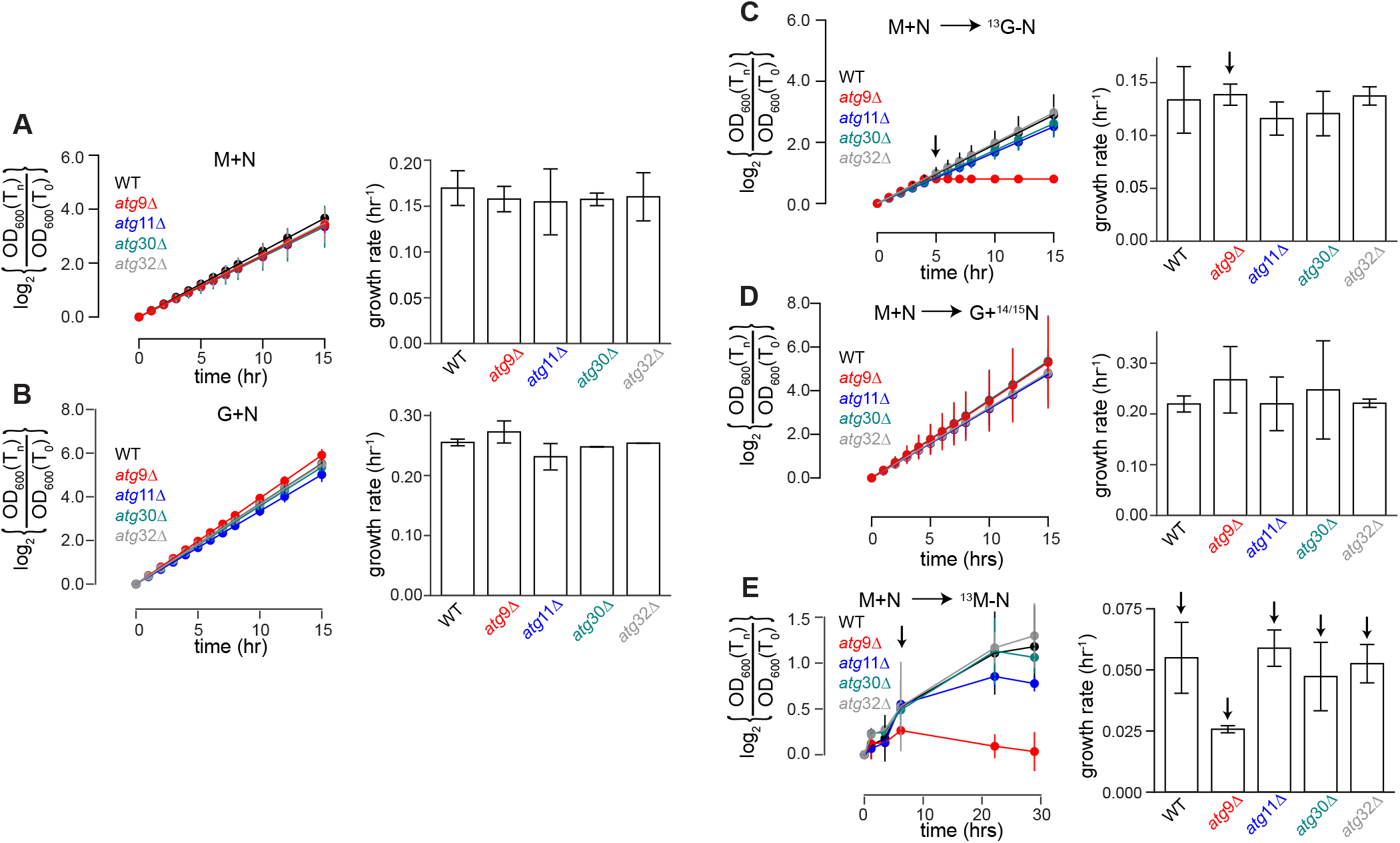
Media-dependent growth of *K. phaffii*. In each growth condition, OD_600_ was monitored in triplicate to assess cell growth. To compare replicates, each OD_600_ measurement was normalized to that at time t=0, and this ratio, after log transformation is plotted (left). Growth rates were calculated by fitting the linear portion of each plot, and a bar chart of the resulting measurements depicts mean ± standard error of the mean (right). For conditions exhibiting non-exponential growth, datapoints ranging from t=0 to that indicated arrow were used to quantify growth rates. **(A)** Steady-state growth in methanol (M+N) media. **(B)** Steady-state growth in glucose (G+N) media. **(C)** At time t=0, cells grown in M+N media were harvested, washed with pre-warmed yeast nitrogen base (YNB), and then resuspended in minimal glucose media lacking nitrogen (G-N). **(D)** Cells were treated as in (C), but were resuspended in glucose media bearing nitrogen (G+N). **(E)** Cells were treated as in (C), but were resuspended in methanol media lacking nitrogen (M-N).

**Supplementary Figure 4.**
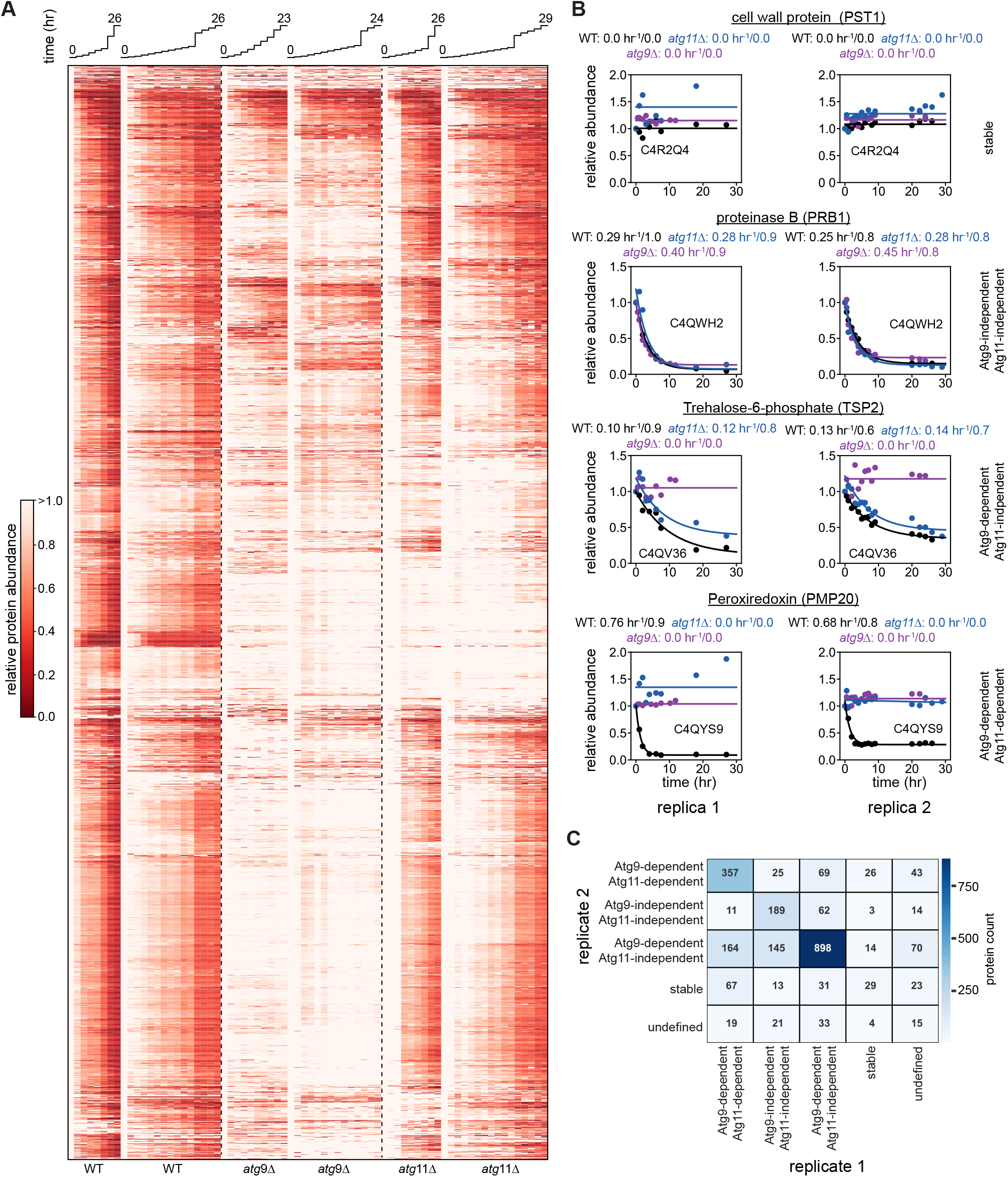
PL-qMS comparison across replicates. **(A)** Clustered heatmap comparing protein degradation as wild-type, *atg*9Δ and *atg*11Δ strains were shifted from growth in methanol (M+N) to glucose media lacking nitrogen (G-N). Columns represent individual timepoints, which are plotted above. **(B)** Measured protein abundance (relative to that at time t=0) plotted against time, with fit (*eq. 1*, see Methods) and fit parameters (*k*_*deg*_/A_deg_) listed. A representative subset of proteins exhibiting different annotated degradation behaviors across listed genetic backgrounds (rows) are highlighted, with two independent replicate PL-qMS trajectories are plotted for each class (columns). **(C)** Confusion matrix comparing the number of proteins assigned to each class between the replicate assays depicted in (A).

**Supplementary Figure 5.**
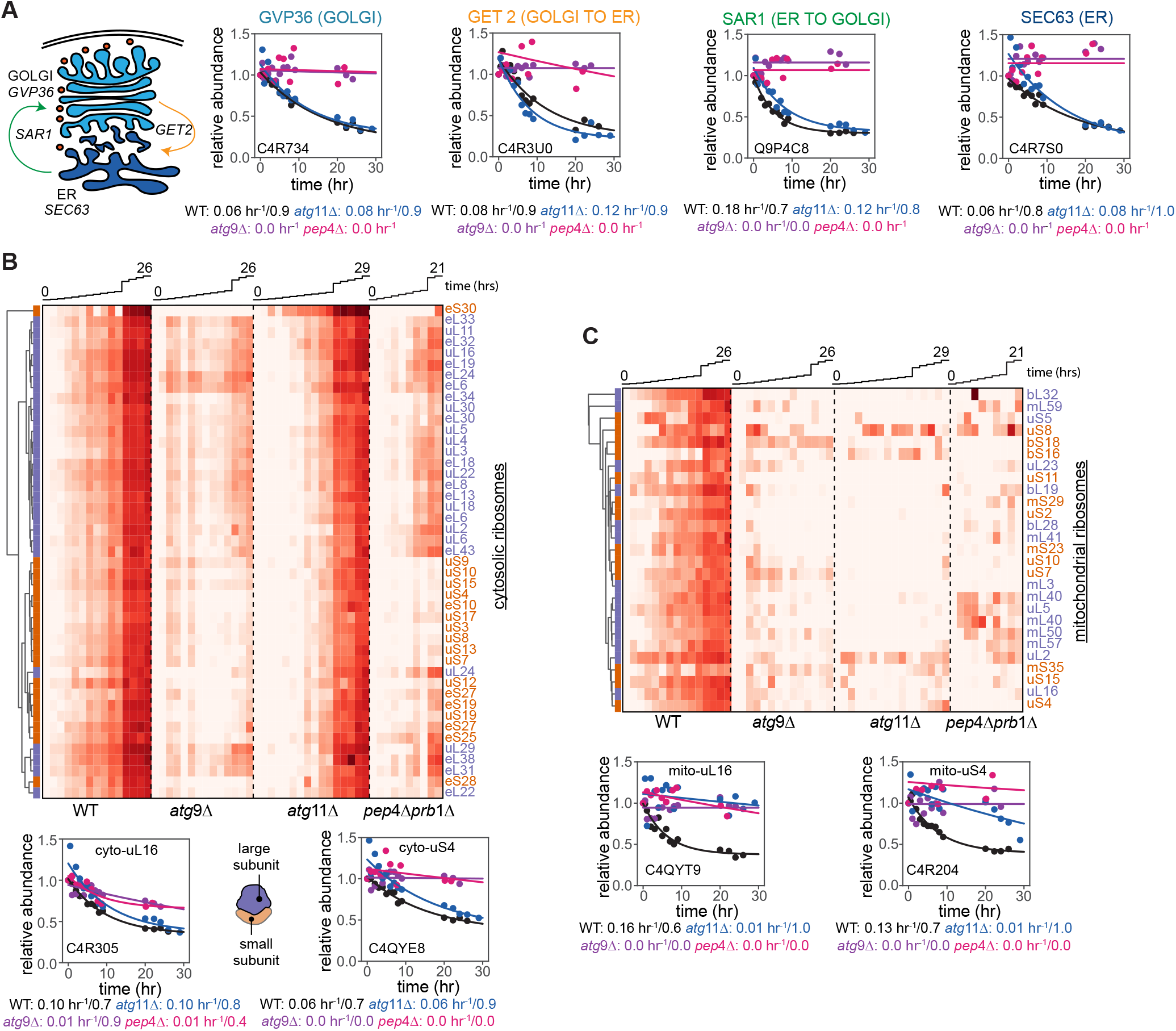
Degradation profiles in cells transitioning from growth on M+N to G-N media for secretory proteins, and for ribosomal, and mito-ribosomal proteins. **(A)** Degradation profiles and fit kinetic models (*eq. 1*, see Methods) for proteins involved in vesicular transport across the ER and Golgi. Fit parameters (*k*_*deg*_/A_deg_) listed, and proteins labels are colored following cartoon of this pathway (top left). Common names and Uniprot IDs noted. **(B)** Clustered heatmap (top) highlighting degradation profile of proteins from the large (blue) and small (orange) cytosolic ribosomal subunits in noted strains. Degradation profiles and fit kinetic models (bottom) for representative large (left) and small (right) cytosolic ribosomal proteins. Note incomplete Atg9- and Pep4/Prb1-dependence for degradation of protein cyto-uL16. **(C)** Clustered heatmap and degradation profiles for mitochondrial ribosomal proteins, as described above. Columns represent individual timepoints, which are plotted above.

**Supplementary Figure 6.**
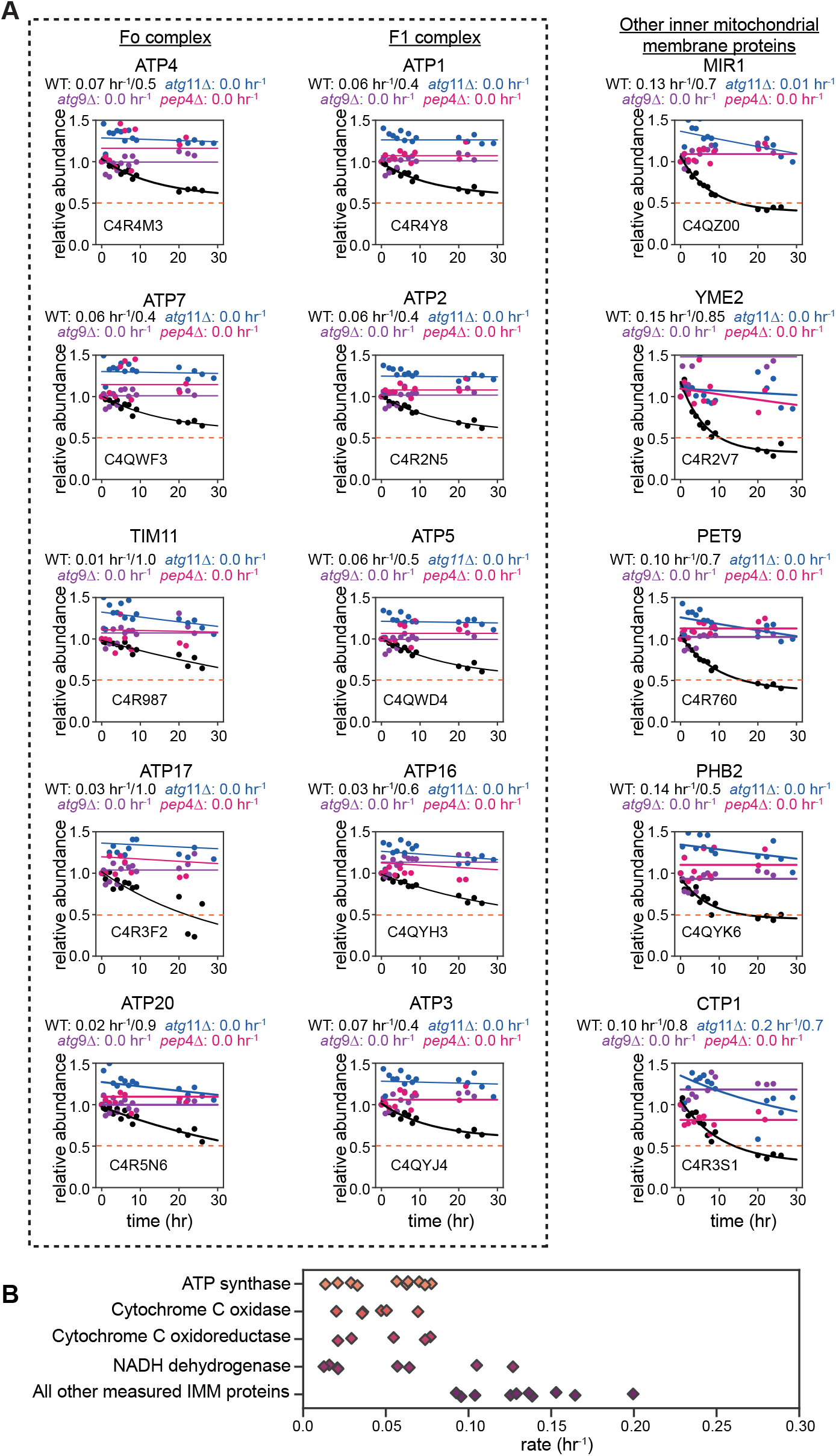
Degradation profiles of mitochondrial proteins in cells transitioning from growth on M+N to G-N media. **(A)** Degradation profiles and fit kinetic models (*eq. 1*, see Methods) for protein subunits of the F_1_/F_o_ ATP synthase complex (left), which were degraded slowly relative to many other inner mitochondrial membrane proteins (right). Dotted line marks 50% degradation. Uniprot ID and common names listed, with fit kinetic parameters (*k*_*deg*_/A_deg_) noted below each fit. **(B)** Fit degradation rates (*k*_*deg*_) for protein subunits of various mitochondrial complexes, with each mark indicating the measured rate for a different subunit. Note the relatively slow degradation the highlighted complexes relative to other inner mitochondrial membrane localized proteins.

**Supplementary Figure 7.**
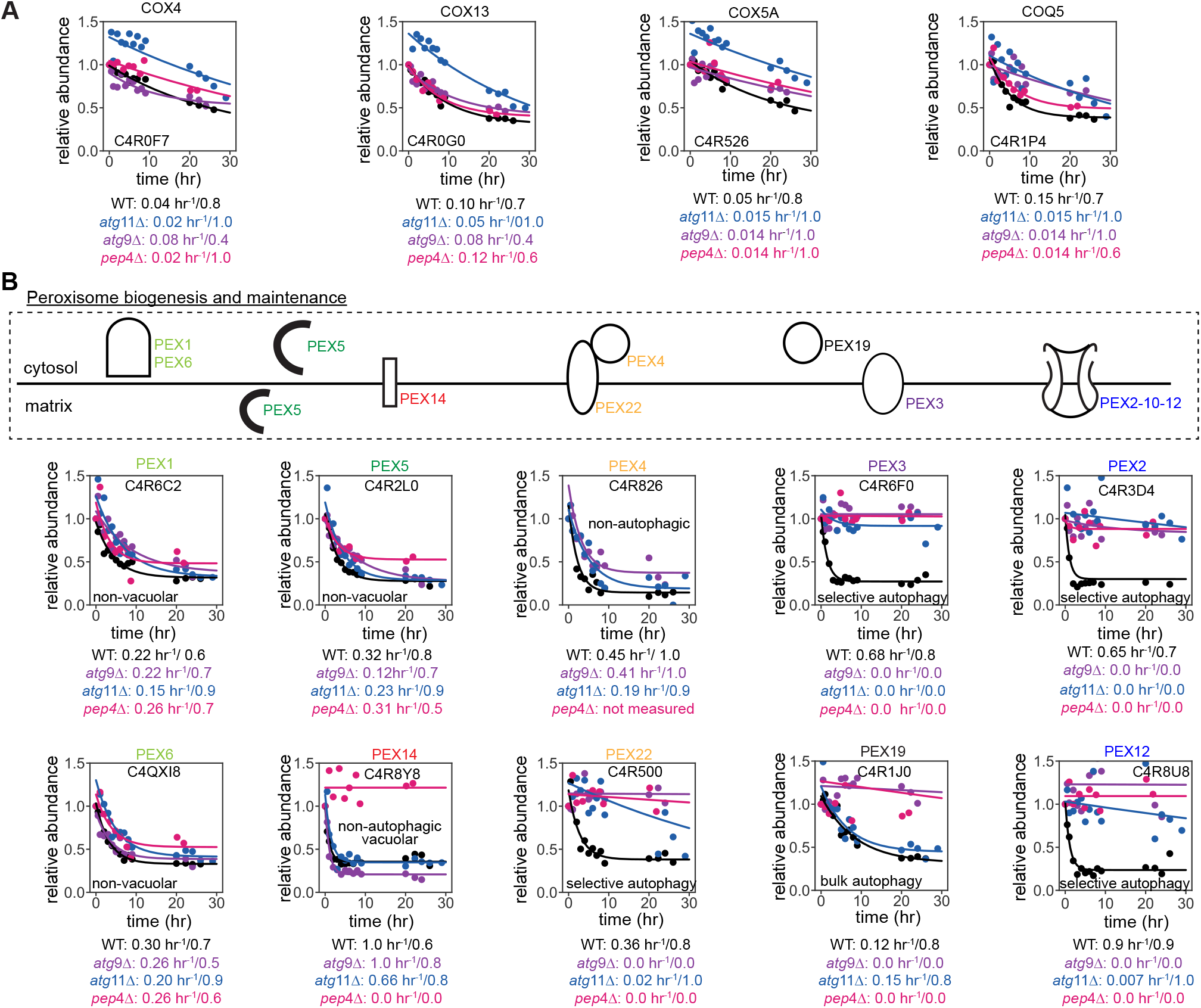
Degradation profiles of selected mitochondrial and peroxisomal proteins in cells transitioning from growth on M+N to G-N media. **(A)** Degradation profiles and fit kinetic models (*eq. 1*, see Methods) for components of the Cox complex, which is localized to the inner mitochondrial membrane. Note that this complex is degraded in an Atg9-, Atg11-, and Pep4/Prb1-independent fashion, in contrast to most mitochondrially localized proteins that were degraded through selective autophagy. Uniprot ID and common names listed, with fit kinetic parameters (*k*_*deg*_/A_deg_) noted below each fit. **(B)** Cartoon of peroxisomal membrane, highlighting localization of a subset of proteins involved in peroxisomal biogenesis and maintenance (top), and degradation profiles and fit kinetic models for these proteins (bottom). Uniprot ID, common name, and assigned mode of degradation listed, with fit kinetic parameters (*k*_*deg*_/A_deg_) noted below each fit.

**Supplementary Figure 8.**
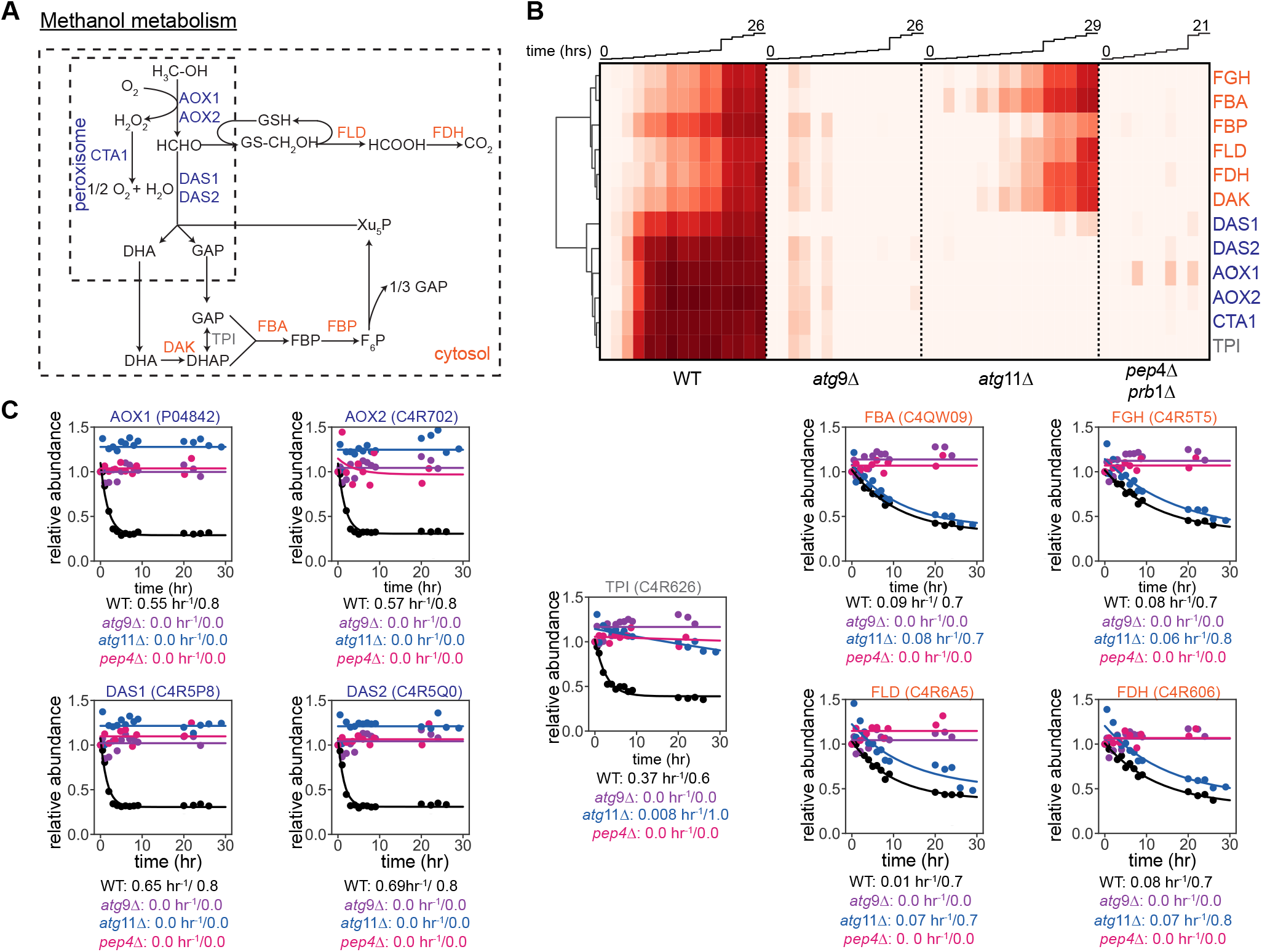
Bulk and selective autophagy collaborate to degraded methanol metabolism related proteins. **(A)** Schematic of methanol metabolism, highlighting cytosolic (orange) and peroxisomal (blue) enzyme localization. Metabolites labeled in black. **(B)** Clustered heatmap highlighting genetic dependence for degradation of proteins involved in methanol utilization as cells transition from growth on M+N to G-N media. **(C)** Degradation profiles and fit kinetic models (*eq. 1*, see Methods) for subset of proteins from panels A-B, separated by localization to the peroxisome (left), cytosol (right). Triose phosphate isomerase is highlighted as its degradation profile is most consistent with peroxisomal degradation, despite being canonically annotated as a protein localized to the cytosol. Uniprot ID, common name, and fit kinetic parameters (*k*_*deg*_/A_deg_) noted with each fit. Columns represent individual timepoints, which are plotted above.

**Supplementary Figure 9.**
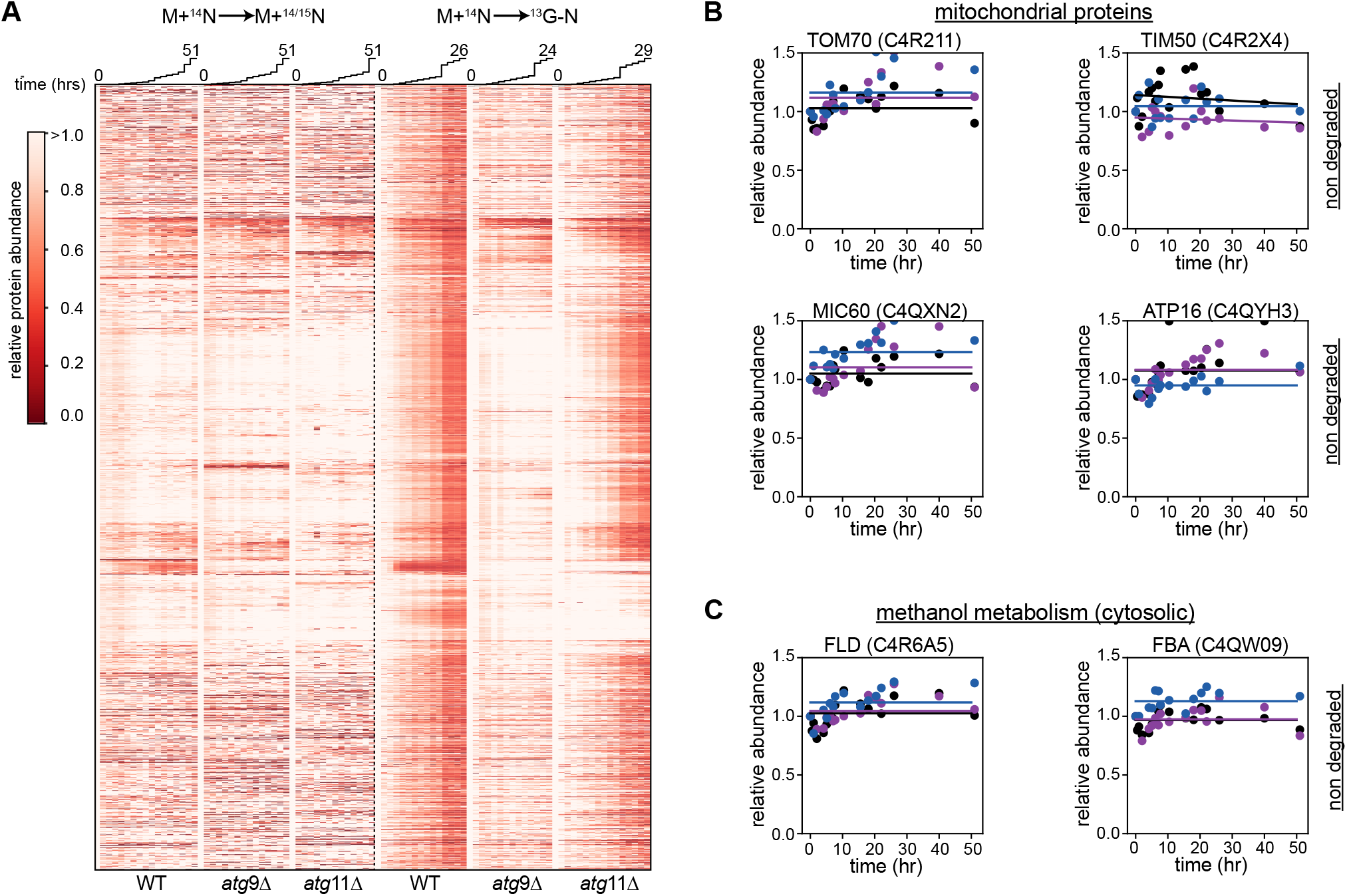
The proteome is largely stable under steady-state growth conditions in M+N media. **(A)** Clustered heatmap comparing proteome-wide degradation in cells grown in steady-state conditions (M+N; left) to that in cells transitioning from M+N to G-N media (right). Columns represent individual timepoints, which are plotted above. **(B)** Degradative profiles for exemplar mitochondrial proteins, which were not degraded in these steady-state M+N media conditions. **(C)** Degradative profiles for exemplar cytosolic proteins involved in methanol utilization. These proteins were not degraded in these steady-state M+N media conditions, whereas those localized to the peroxisome were degraded in an Atg9- and Atg11-dependent fashion (see Figure 3).

**Supplementary Figure 10.**
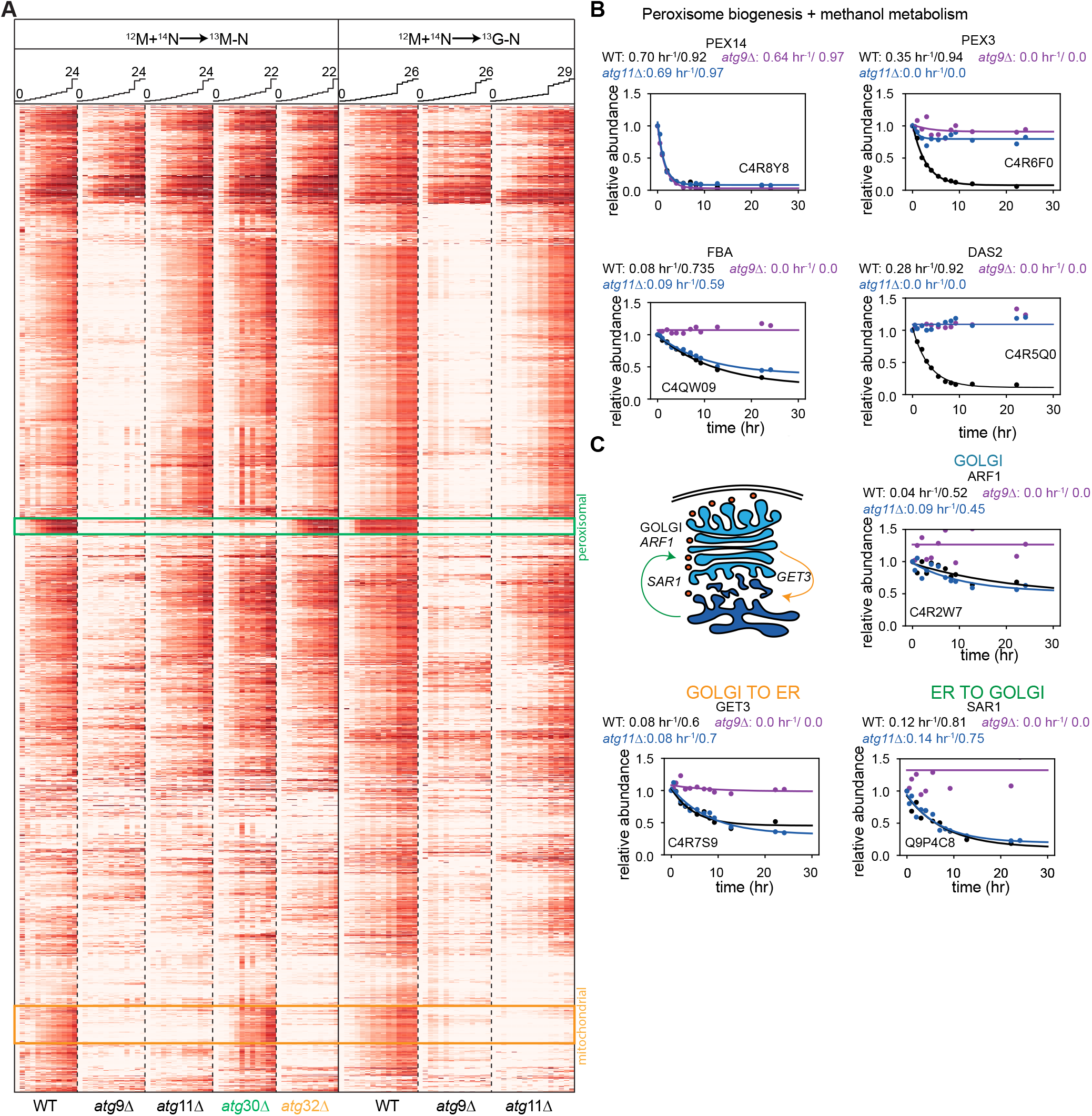
Nitrogen starvation in methanol and glucose promote similar proteome remodeling responses. **(A)** Clustered heatmap comparing proteome remodeling as cells transition from M+^14^N to ^13^M-N versus M+^14^N to ^13^G-N in various genetic backgrounds. Columns represent individual timepoints, which are plotted above. Peroxisomal and mitochondrial proteins highlighted in green and orange boxes, respectively. **(B)** The degradation profiles of proteins involved in peroxisome biogenesis and methanol metabolism during cellular adaptation from M+^14^N to ^13^G-N media. **(C)** Degradation profiles and fit kinetic models (*eq. 1*, see Methods) of proteins involved in vesicular transport during cellular adaptation from M+^14^N to ^13^G-N media. Uniprot ID, common name, and fit kinetic parameters (*k*_*deg*_/A_deg_) noted with each fit.

## SUPPLEMENTARY TABLES

**Supplementary Table 1:** Cell-wall localized, degradation-resistant proteins. Columns list Uniprot ID, common name, and description for stable proteins annotated to localize to the cell-wall (Burgard *et al*., 2020). Proteins lacking an annotated common name listed as n/a.

| Uniprot ID | Common name | Description (Uniprot taxonomy description field) |
| --- | --- | --- |
| C4QYF3 | BGL2 | Endo-beta-1,3-glucanase, major protein of the cell wall, involved in cell wall maintenance |
| C4QW71 | DSE4 | Daughter cell-specific secreted protein with similarity to glucanases, endo-1,3-beta-glucanase |
| C4R3H3 | EPX1 | SCP domain-containing protein |
| C4R0Q7 | EXG1 | Major exo-1,3-beta-glucanase of the cell wall, involved in cell wall beta-glucan assembly |
| C4QVL4 | GAS1-1 | 1,3-beta-glucanosyltransferase |
| C4R9F4 | GAS3 | 1,3-beta-glucanosyltransferase |
| C4R0Z8 | n/a | Uncharacterized protein with predicted GPI-attachment site |
| C4R3C4 | n/a | Uncharacterized protein with predicted GPI-attachment site |
| C4R2Q4 | PST1 | Cell wall protein that contains a putative GPI-attachment site |
| C4QVL7 | SCW10 | Cell wall protein with similarity to glucanases |
| C4QZH9 | SCW11 | Cell wall protein with similarity to glucanases |
| C4R6P9 | UTH1 | Mitochondrial outer membrane and cell wall localized SUN family member |
| C4QWP6 | UTR2 | Glycosidase |
| C4R8B8 | YPS1-1 | Aspartic protease, attached to the plasma membrane via a glycosylphosphatidylinositol (GPI) anchor |

**Supplementary Table 2:** Proteins degraded in a Atg9-independent, Prb4/Prb1-dependent fashion. Columns list Uniprot ID, common name, and description. Proteins lacking an annotated common name listed as n/a.

| Uniprot ID | Common name | Description (Uniprot taxonomy description field) |
| --- | --- | --- |
| C4QV09 | n/a | DUF221-domain-containing protein |
| C4QV32 | BIO2 | Biotin synthase, catalyzes the conversion of dethiobiotin to biotin |
| C4QVF4 | n/a | Uncharacterized protein |
| C4R522 | SRP1 | Importin subunit alpha |
| C4QW23 | HLJ1 | Co-chaperone for Hsp40p, anchored in the ER membrane |
| C4QWB6 | PIM1-2 | Lon protease homolog 2, peroxisomal |
| C4QWE1 | AYF2 | Arylsulfotransferase |
| C4QWH2 | PRB1 | Vacuolar proteinase B (YscB), a serine protease of the subtilisin family |
| C4QWW6 | UFD4 | Ubiquitin-protein ligase (E3) that interacts with Rpt4p and Rpt6p |
| C4QWZ7 | CHS1 | Chitin synthase |
| C4QX33 | ENA2 | P-type ATPase sodium pump, involved in Na<sup>+</sup> and Li<sup>+</sup> efflux to allow salt tolerance |
| C4QX53 | PST2 | Protein with similarity to members of a family of flavodoxin-like proteins |
| C4QY57 | ICL1 | Isocitrate lyase |
| C4QZ79 | TNA1-2 | Vitamin H transporter 1 |
| C4QZA6 | ROT1 | Protein ROT1 |
| C4QZH7 | STE6 | Plasma membrane ATP-binding cassette (ABC) transporter required for the export of a-factor |
| C4R075 | FZO1 | Mitochondrial integral membrane protein involved in mitochondrial fusion and maintenance of the mitochondria |
| C4R0D6 | n/a | DUF3844 domain-containing protein |
| C4R0L9 | EPS1 | ER protein with chaperone and co-chaperone activity, involved in retention of resident ER proteins |
| C4R0Y0 | BUD22 | Mitochondrial 3-oxoacyl-[acyl-carrier-protein] reductase |
| C4R147 | GRX6 | Cis-golgi localized monothiol glutaredoxin that binds an iron-sulfur cluster |
| C4R1D2 | FKS1 | Catalytic subunit of 1,3-beta-D-glucan synthase |
| C4R1W5 | VTC4 | Vacuolar membrane protein involved in vacuolar polyphosphate accumulation |
| C4R2D2 | n/a | Uncharacterized protein |
| C4R2S3 | NSA2 | Protein constituent of 66S pre-ribosomal particles, contributes to processing of the 27S pre-rRNA |
| C4R3T3 | PEX25 | Peripheral peroxisomal membrane peroxin |
| C4R3V8 | KRE2 | Alpha1,2-mannosyltransferase |
| C4R4D0 | CCC1 | Putative vacuolar Fe<sup>2+</sup>/Mn<sup>2+</sup> transporter |
| C4R560 | THI4 | Thiamine thiazole synthase |
| C4R5Q9 | PDR17 | Phosphatidylinositol transfer protein (PITP) |
| C4R6R4 | ERG4 | C-24(28) sterol reductase, catalyzes the final step in ergosterol biosynthesis |
| C4R7V4 | ORM1 | Evolutionarily conserved protein with similarity to Orm2p |
| C4R7Z9 | CXF1 | Conserved protein involved in exocytic transport from the Golgi |
| C4R8Y8 | PEX14 | Peroxisomal membrane peroxin that is a central component of the peroxisomal protein import machinery |
| C4R915 | COQ1 | Hexaprenyl pyrophosphate synthetase |
| C4R922 | ITR100 | Myo-inositol transporter with strong similarity to the minor myo-inositol transporter Itr |
| C4R6W2 | GLY1 | Threonine aldolase |
| C4QV19 | PAM16 | Constituent of the mitochondrial import motor associated with the presequence translocase |
| C4QZ04 | YCF1 | Metal resistance protein YCF1 |
| C4R070 | SNQ2 | Plasma membrane ATP-binding cassette (ABC) transporter, multidrug transporter involved in multidrug |
| C4R0F3 | YCK3 | Palmitoylated, plasma membrane-bound casein kinase I isoform |
| C4R0Y6 | n/a | Protein FMP42 |
| C4R1C0 | n/a | Uncharacterized protein |
| C4R1E0 | VTC2 | Vacuolar membrane protein involved in vacuolar polyphosphate accumulation |
| C4R4D6 | OYE3 | Widely conserved NADPH oxidoreductase containing flavin mononucleotide (FMN) |
| C4R4H7 | CMP2 | Serine/threonine-protein phosphatase |
| C4R846 | ZRT2 | Low-affinity zinc transporter of the plasma membrane |

**Supplementary Table 3:**
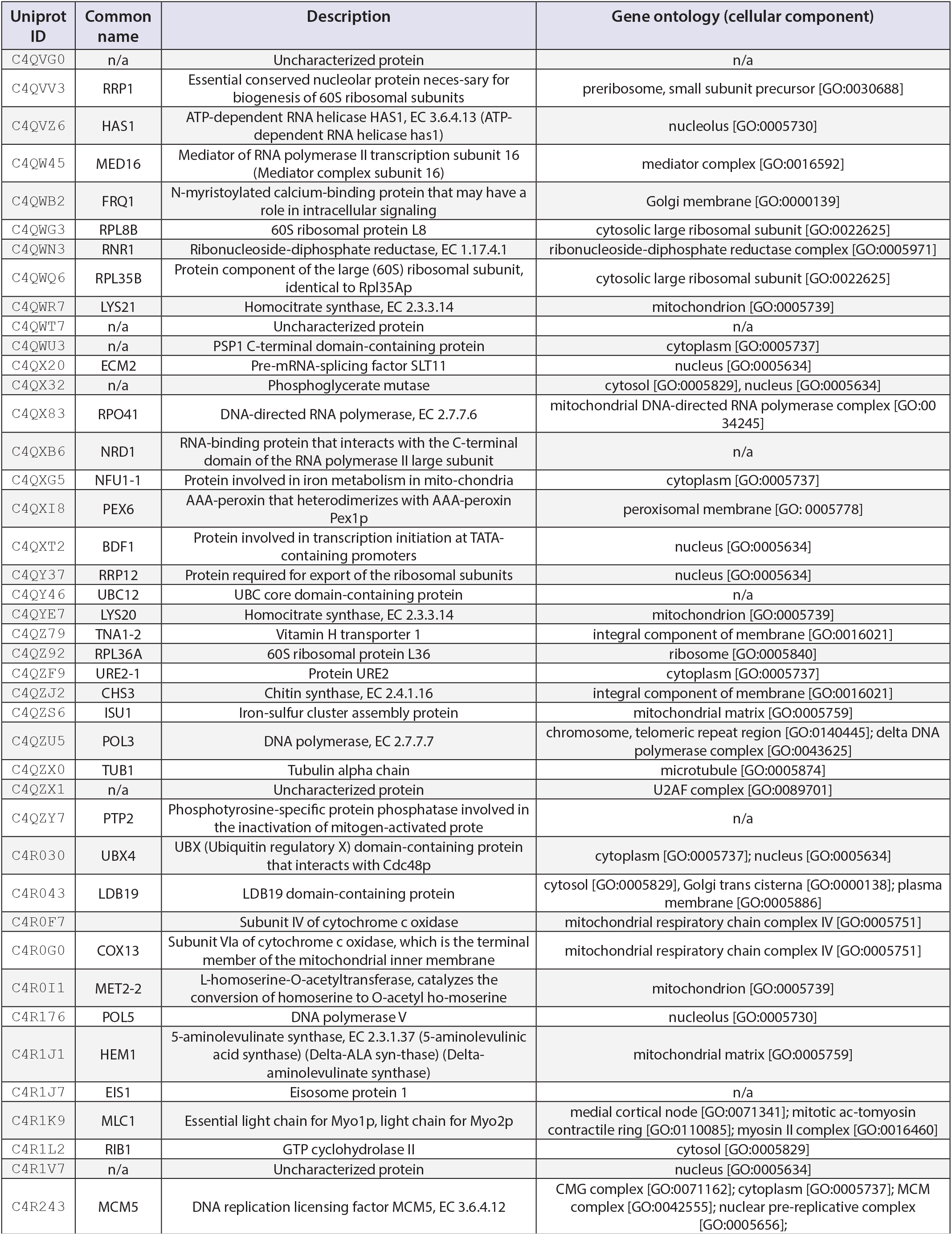
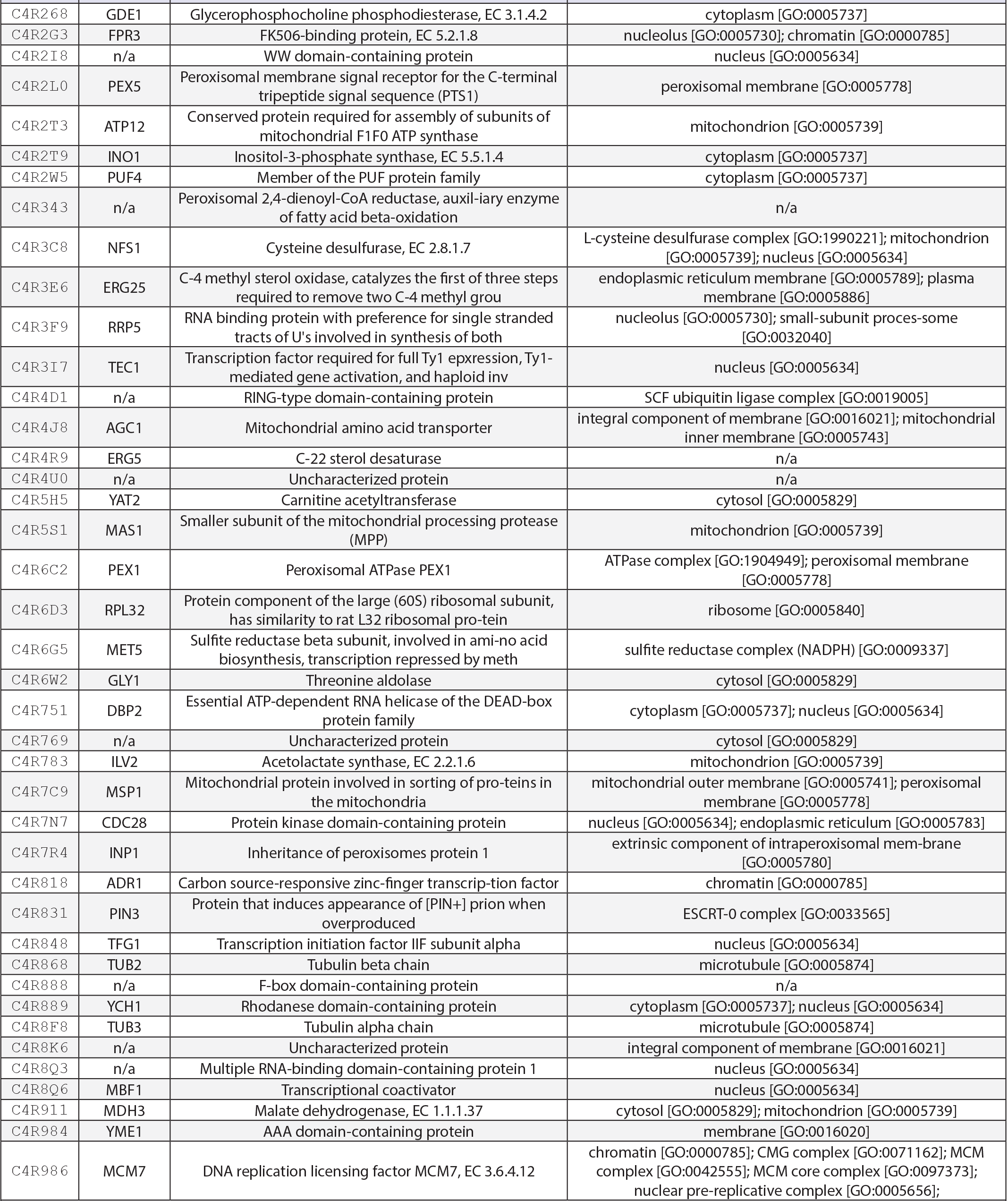
Proteins degraded in Prb4/Prb1-independent fashion. Columns list Uniprot ID, common name, description, and associated gene ontology annotations. Proteins lacking an annotated common name listed as n/a. Table continued on following page.

| Uniprot ID | Common name | Description | Gene ontology (cellular component) |
| --- | --- | --- | --- |
| C4QVG0 | n/a | Uncharacterized protein | n/a |
| C4QVV3 | RRP1 | Essential conserved nucleolar protein necessary for biogenesis of 60S ribosomal subunits | preribosome, small subunit precursor [GO:0030688] |
| C4QVZ6 | HAS1 | ATP-dependent RNA helicase HAS1, EC 3.6.4.13 (ATP-dependent RNA helicase has1) | nucleolus [GO:0005730] |
| C4QW45 | MED16 | Mediator of RNA polymerase II transcription subunit 16 (Mediator complex subunit 16) | mediator complex [GO:0016592] |
| C4QWB2 | FRQ1 | N-myristoylated calcium-binding protein that may have a role in intracellular signaling | Golgi membrane [GO:0000139] |
| C4QWG3 | RPL8B | 60S ribosomal protein L8 | cytosolic large ribosomal subunit [GO:0022625] |
| C4QWN3 | RNR1 | Ribonucleoside-diphosphate reductase, EC 1.17.4.1 | ribonucleoside-diphosphate reductase complex [GO:0005971] |
| C4QWQ6 | RPL35B | Protein component of the large (60S) ribosomal subunit, identical to Rpl35Ap | cytosolic large ribosomal subunit [GO:0022625] |
| C4QWR7 | LYS21 | Homocitrate synthase, EC 2.3.3.14 | mitochondrion [GO:0005739] |
| C4QWT7 | n/a | Uncharacterized protein | n/a |
| C4QWU3 | n/a | PSP1 C-terminal domain-containing protein | cytoplasm [GO:0005737] |
| C4QX20 | ECM2 | Pre-mRNA-splicing factor SLT11 | nucleus [GO:0005634] |
| C4QX32 | n/a | Phosphoglycerate mutase | cytosol [GO:0005829], nucleus [GO:0005634] |
| C4QX83 | RPO41 | DNA-directed RNA polymerase, EC 2.7.7.6 | mitochondrial DNA-directed RNA polymerase complex [GO:0034245] |
| C4QXB6 | NRD1 | RNA-binding protein that interacts with the C-terminal domain of the RNA polymerase II large subunit | n/a |
| C4QXG5 | NFU1-1 | Protein involved in iron metabolism in mitochondria | cytoplasm [GO:0005737] |
| C4QXI8 | PEX6 | AAA-peroxin that heterodimerizes with AAA-peroxin Pex1p | peroxisomal membrane [GO:0005778] |
| C4QXT2 | BDF1 | Protein involved in transcription initiation at TATA-containing promoters | nucleus [GO:0005634] |
| C4QY37 | RRP12 | Protein required for export of the ribosomal subunits | nucleus [GO:0005634] |
| C4QY46 | UBC12 | UBC core domain-containing protein | n/a |
| C4QYE7 | LYS20 | Homocitrate synthase, EC 2.3.3.14 | mitochondrion [GO:0005739] |
| C4QZ79 | TNA1-2 | Vitamin H transporter 1 | integral component of membrane [GO:0016021] |
| C4QZ92 | RPL36A | 60S ribosomal protein L36 | ribosome [GO:0005840] |
| C4QZF9 | URE2-1 | Protein URE2 | cytoplasm [GO:0005737] |
| C4QZJ2 | CHS3 | Chitin synthase, EC 2.4.1.16 | integral component of membrane [GO:0016021] |
| C4QZS6 | ISU1 | Iron-sulfur cluster assembly protein | mitochondrial matrix [GO:0005759] |
| C4QZU5 | POL3 | DNA polymerase, EC 2.7.7.7 | chromosome, telomeric repeat region [GO:0140445]; delta DNA polymerase complex [GO:0043625] |
| C4QZX0 | TUB1 | Tubulin alpha chain | microtubule [GO:0005874] |
| C4QZX1 | n/a | Uncharacterized protein | U2AF complex [GO:0089701] |
| C4QZY7 | PTP2 | Phosphotyrosine-specific protein phosphatase involved in the inactivation of mitogen-activated prote | n/a |
| C4R030 | UBX4 | UBX (Ubiquitin regulatory X) domain-containing protein that interacts with Cdc48p | cytoplasm [GO:0005737]; nucleus [GO:0005634] |
| C4R043 | LDB19 | LDB19 domain-containing protein | cytosol [GO:0005829], Golgi trans cisterna [GO:0000138]; plasma membrane [GO:0005886] |
| C4R0F7 |  | Subunit IV of cytochrome c oxidase | mitochondrial respiratory chain complex IV [GO:0005751] |
| C4R0G0 | COX13 | Subunit VIa of cytochrome c oxidase, which is the terminal member of the mitochondrial inner membrane | mitochondrial respiratory chain complex IV [GO:0005751] |
| C4R0I1 | MET2-2 | L-homoserine-O-acetyltransferase, catalyzes the conversion of homoserine to O-acetyl homoserine | mitochondrion [GO:0005739] |
| C4R176 | POL5 | DNA polymerase V | nucleolus [GO:0005730] |
| C4R1J1 | HEM1 | 5-aminolevulinate synthase, EC 2.3.1.37 (5-aminolevulinic acid synthase) (Delta-ALA synthase) (Delta-aminolevulinate synthase) | mitochondrial matrix [GO:0005759] |
| C4R1J7 | EIS1 | Eisosome protein 1 | n/a |
| C4R1K9 | MLC1 | Essential light chain for Myo1p, light chain for Myo2p | medial cortical node [GO:0071341]; mitotic actomyosin contractile ring [GO:0110085]; myosin II complex [GO:0016460] |
| C4R1L2 | RIB1 | GTP cyclohydrolase II | cytosol [GO:0005829] |
| C4R1V7 | n/a | Uncharacterized protein | nucleus [GO:0005634] |
| C4R243 | MCM5 | DNA replication licensing factor MCM5, EC 3.6.4.12 | CMG complex [GO:0071162]; cytoplasm [GO:0005737]; MCM complex [GO:0042555]; nuclear pre-replicative complex [GO:0005656]; |
| C4R268 | GDE1 | Glycerophosphocholine phosphodiesterase, EC 3.1.4.2 | cytoplasm [GO:0005737] |
| C4R2G3 | FPR3 | FK506-binding protein, EC 5.2.1.8 | nucleolus [GO:0005730]; chromatin [GO:0000785] |
| C4R2I8 | n/a | WW domain-containing protein | nucleus [GO:0005634] |
| C4R2L0 | PEX5 | Peroxisomal membrane signal receptor for the C-terminal tripeptide signal sequence (PTS1) | peroxisomal membrane [GO:0005778] |
| C4R2T3 | ATP12 | Conserved protein required for assembly of subunits of mitochondrial F1F0 ATP synthase | mitochondrion [GO:0005739] |
| C4R2T9 | INO1 | Inositol-3-phosphate synthase, EC 5.5.1.4 | cytoplasm [GO:0005737] |
| C4R2W5 | PUF4 | Member of the PUF protein family | cytoplasm [GO:0005737] |
| C4R343 | n/a | Peroxisomal 2,4-dienoyl-CoA reductase, auxil-iary enzyme of fatty acid beta-oxidation | n/a |
| C4R3C8 | NFS1 | Cysteine desulfurase, EC 2.8.1.7 | L-cysteine desulfurase complex [GO:1990221]; mitochondrion [GO:0005739]; nucleus [GO:0005634] |
| C4R3E6 | ERG25 | C-4 methyl sterol oxidase, catalyzes the first of three steps required to remove two C-4 methyl grou | endoplasmic reticulum membrane [GO:0005789]; plasma membrane [GO:0005886] |
| C4R3F9 | RRP5 | RNA binding protein with preference for single stranded tracts of U's involved in synthesis of both | nucleolus [GO:0005730]; small-subunit proces-some [GO:0032040] |
| C4R3I7 | TEC1 | Transcription factor required for full Ty1 epxression, Ty1-mediated gene activation, and haploid inv | nucleus [GO:0005634] |
| C4R4D1 | n/a | RING-type domain-containing protein | SCF ubiquitin ligase complex [GO:0019005] |
| C4R4J8 | AGC1 | Mitochondrial amino acid transporter | integral component of membrane [GO:0016021]; mitochondrial inner membrane [GO:0005743] |
| C4R4R9 | ERG5 | C-22 sterol desaturase | n/a |
| C4R4U0 | n/a | Uncharacterized protein | n/a |
| C4R5H5 | YAT2 | Carnitine acetyltransferase | cytosol [GO:0005829] |
| C4R5S1 | MAS1 | Smaller subunit of the mitochondrial processing protease (MPP) | mitochondrion [GO:0005739] |
| C4R6C2 | PEX1 | Peroxisomal ATPase PEX1 | ATPase complex [GO:1904949]; peroxisomal membrane [GO:0005778] |
| C4R6D3 | RPL32 | Protein component of the large (60S) ribosomal subunit, has similarity to rat L32 ribosomal pro-tein | ribosome [GO:0005840] |
| C4R6G5 | MET5 | Sulfite reductase beta subunit, involved in ami-no acid biosynthesis, transcription repressed by meth | sulfite reductase complex (NADPH) [GO:0009337] |
| C4R6W2 | GLY1 | Threonine aldolase | cytosol [GO:0005829] |
| C4R751 | DBP2 | Essential ATP-dependent RNA helicase of the DEAD-box protein family | cytoplasm [GO:0005737]; nucleus [GO:0005634] |
| C4R769 | n/a | Uncharacterized protein | cytosol [GO:0005829] |
| C4R783 | ILV2 | Acetolactate synthase, EC 2.2.1.6 | mitochondrion [GO:0005739] |
| C4R7C9 | MSP1 | Mitochondrial protein involved in sorting of pro-teins in the mitochondria | mitochondrial outer membrane [GO:0005741]; peroxisomal membrane [GO:0005778] |
| C4R7N7 | CDC28 | Protein kinase domain-containing protein | nucleus [GO:0005634]; endoplasmic reticulum [GO:0005783] |
| C4R7R4 | INP1 | Inheritance of peroxisomes protein 1 | extrinsic component of intraperoxisomal mem-brane [GO:0005780] |
| C4R818 | ADR1 | Carbon source-responsive zinc-finger transcrip-tion factor | chromatin [GO:0000785] |
| C4R831 | PIN3 | Protein that induces appearance of [PIN+] prion when overproduced | ESCRT-0 complex [GO:0033565] |
| C4R848 | TFG1 | Transcription initiation factor IIF subunit alpha | nucleus [GO:0005634] |
| C4R868 | TUB2 | Tubulin beta chain | microtubule [GO:0005874] |
| C4R888 | n/a | F-box domain-containing protein | n/a |
| C4R889 | YCH1 | Rhodanese domain-containing protein | cytoplasm [GO:0005737]; nucleus [GO:0005634] |
| C4R8F8 | TUB3 | Tubulin alpha chain | microtubule [GO:0005874] |
| C4R8K6 | n/a | Uncharacterized protein | integral component of membrane [GO:0016021] |
| C4R8Q3 | n/a | Multiple RNA-binding domain-containing protein 1 | nucleus [GO:0005634] |
| C4R8Q6 | MBF1 | Transcriptional coactivator | nucleus [GO:0005634] |
| C4R911 | MDH3 | Malate dehydrogenase, EC 1.1.1.37 | cytosol [GO:0005829]; mitochondrion [GO:0005739] |
| C4R984 | YME1 | AAA domain-containing protein | membrane [GO:0016020] |
| C4R986 | MCM7 | DNA replication licensing factor MCM7, EC 3.6.4.12 | chromatin [GO:0000785]; CMG complex [GO:0071162]; MCM complex [GO:0042555]; MCM core complex [GO:0097373]; nuclear pre-replicative complex [GO:0005656]; |
| **Supplementary Table 3 (continued): Proteins degraded in Prb4/Prb1-independent fashion.** Columns list Uniprot ID, common name, description, and associated gene ontology annotations. Proteins lacking an annotated common name listed as n/a. |  |  |  |

**Supplementary Table 4:**
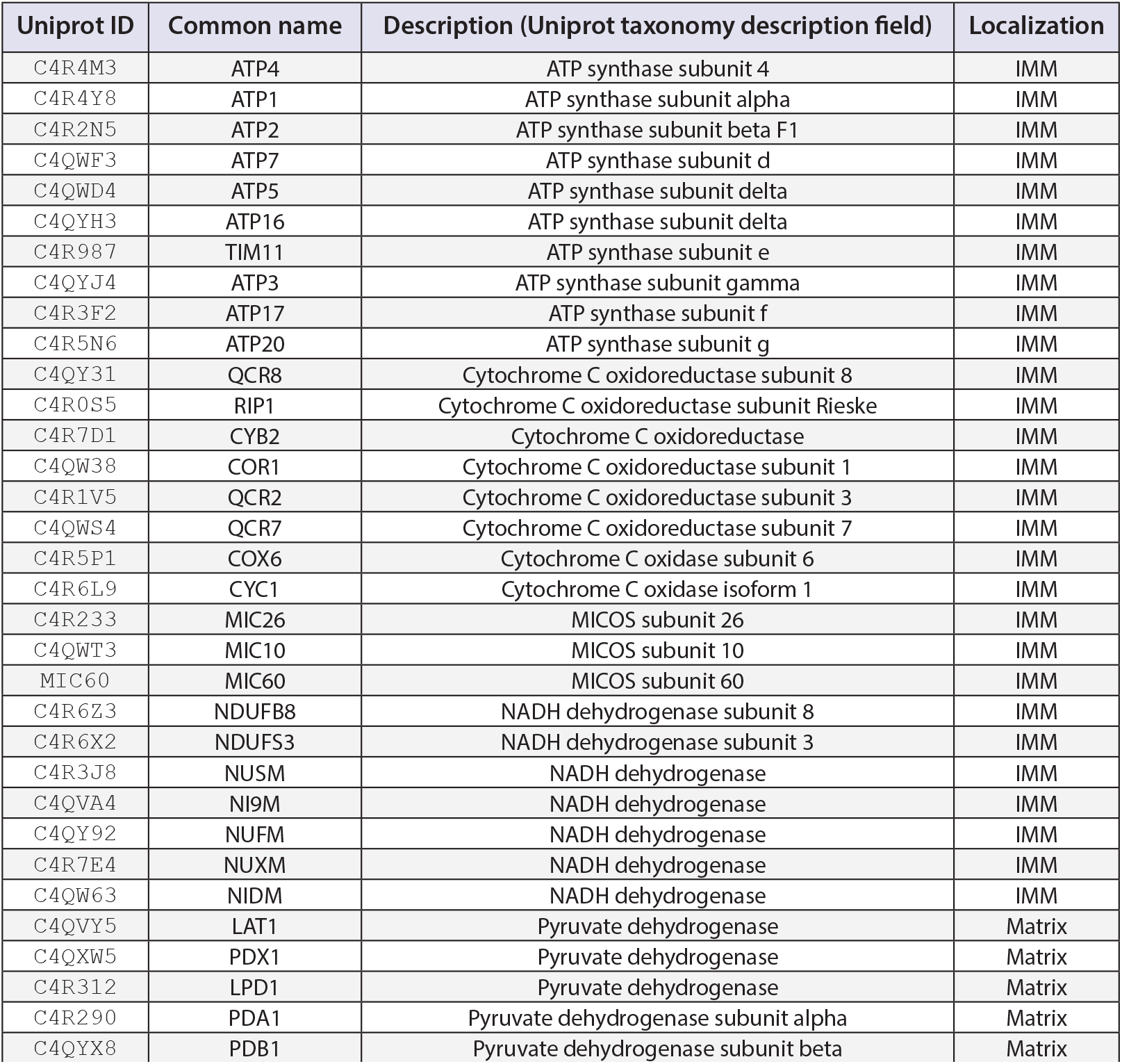
Mitochondrial proteins exhibiting relatively slow degradation kinetics. Full list of mitochondrially localized proteins highlighted for their slow degradation kinetics (see Figure 2B). Columns list Uniprot ID, common name, description, and localization within the mitochondria.

**Supplementary Table 5:** Degradative mode of proteins involved in peroxisome biogenesis and methanol metabolism. Columns list Uniprot ID, common name, assigned mode of degradation, and pathway.

| Uniprot ID | Common name | Mode of degradation | Pathway |
| --- | --- | --- | --- |
| C4R6C2 | PEX1 | Atg9-Atg11-independent, vacuole-independent | peroxisome biogenesis |
| C4R2L0 | PEX5 | Atg9-Atg11-independent, vacuole-independent |  |
| C4QXI8 | PEX6 | Atg9-Atg11-independent, vacuole-independent |  |
| C4R826 | PEX4 | Atg9-Atg11-independent |  |
| C4R8Y8 | PEX14 | Atg9-Atg11-independent, vacuole-dependent |  |
| C4R3T3 | PEX25 | Atg9-Atg11-independent, vacuole-dependent |  |
| C4R825 | PEX26 | Atg9-Atg11-independent, vacuole-dependent |  |
| C4R3D4 | PEX2 | Atg9-Atg11-dependent, vacuole-dependent |  |
| C4R6F0 | PEX3 | Atg9-Atg11-dependent, vacuole-dependent |  |
| C4QY68 | PEX8 | Atg9-Atg11-dependent, vacuole-dependent |  |
| C4R0W1 | PEX11 | Atg9-Atg11-dependent, vacuole-dependent |  |
| C4R8U8 | PEX12 | Atg9-Atg11-dependent, vacuole-dependent |  |
| C4R500 | PEX22 | Atg9-Atg11-dependent, vacuole-dependent |  |
| C4R1J0 | PEX19 | Atg9-dependent, vacuole-dependent |  |
| C4QYD8 | PEX30 | Atg9-dependent, vacuole-dependent |  |
| ALOX1 | AOX1 | Atg9-Atg11-dependent, vacuole-dependent | methanol metabolism |
| ALOX2 | AOX2 | Atg9-Atg11-dependent, vacuole-dependent |  |
| C4R2S1 | CTA1 | Atg9-Atg11-dependent, vacuole-dependent |  |
| C4R5P8 | DAS1 | Atg9-Atg11-dependent, vacuole-dependent |  |
| C4R5Q0 | DAS2 | Atg9-Atg11-dependent, vacuole-dependent |  |
| C4R6A5 | FLD | Atg9-dependent, vacuole-dependent |  |
| C4R606 | FDH | Atg9-dependent, vacuole-dependent |  |
| C4R5Q6 | DAK | Atg9-dependent, vacuole-dependent |  |
| C4R5T5 | FGH | Atg9-dependent, vacuole-dependent |  |
| C4R626 | TPI | Atg9-dependent, vacuole-dependent |  |
| C4R5T8 | FBP | Atg9-dependent, vacuole-dependent |  |
| C4QW09 | FBA | Atg9-dependent, vacuole-dependent |  |

**Supplementary Table 6:** *Komagataella phaffii* strains used in this study. Columns detail strain name within the Davis lab repository, description as used in this study, genotype; and strain source.

| Strain name | Description | Genotype | Source |
| --- | --- | --- | --- |
| st_JD714 | wild-type | GS115 pIB3::HIS4 | Subramani lab |
| st_JD715 | *atg9Δ* | GS115 *atg9Δ*:: Zeocin<sup>R</sup> pIB::HIS4 | Subramani lab |
| st_JD716 | *atg11Δ* | GS115 *atg11Δ*:: Zeocin<sup>R</sup> pIB::HIS4 | Subramani lab |
| st_JD719 | *atg30Δ* | GS115 *atg30Δ*:: Zeocin<sup>R</sup> pIB::HIS4 | Subramani lab |
| st_JD720 | *atg32Δ* | GS115 *atg32Δ*:: Zeocin<sup>R</sup> pIB::HIS4 | Subramani lab |
| st_JD716 | *pep4Δprb1Δ* | SMD1163 *pep4Δ/prb1Δ/his4Δ* pIB3::HIS4 | This work |

**Supplementary Table 7:**
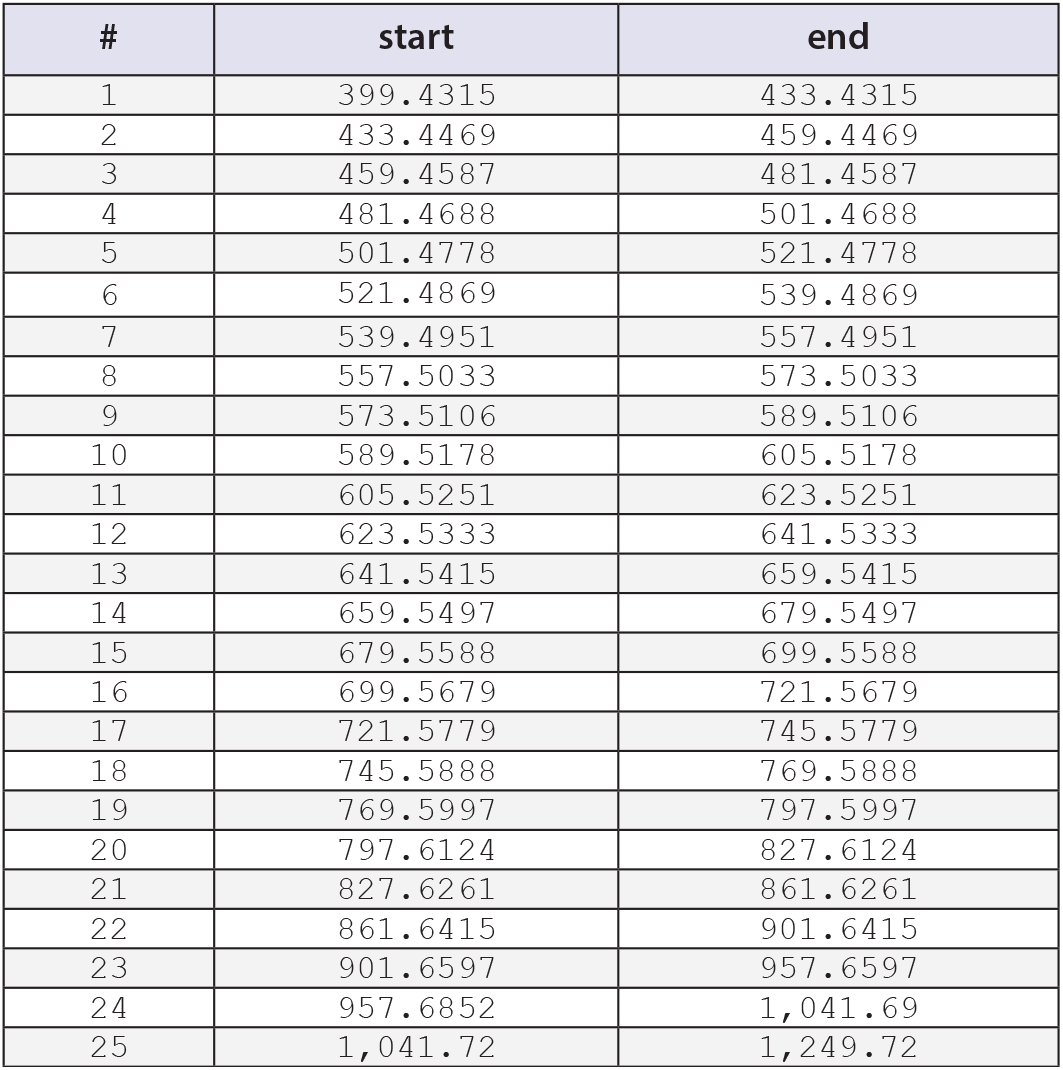
Data-independent aquisition (DIA) windows. Start and end m/z values for DIA acquisition windows used in this study.

| # | start | end |
| --- | --- | --- |
| 1 | 399.4315 | 433.4315 |
| 2 | 433.4469 | 459.4469 |
| 3 | 459.4587 | 481.4587 |
| 4 | 481.4688 | 501.4688 |
| 5 | 501.4778 | 521.4778 |
| 6 | 521.4869 | 539.4869 |
| 7 | 539.4951 | 557.4951 |
| 8 | 557.5033 | 573.5033 |
| 9 | 573.5106 | 589.5106 |
| 10 | 589.5178 | 605.5178 |
| 11 | 605.5251 | 623.5251 |
| 12 | 623.5333 | 641.5333 |
| 13 | 641.5415 | 659.5415 |
| 14 | 659.5497 | 679.5497 |
| 15 | 679.5588 | 699.5588 |
| 16 | 699.5679 | 721.5679 |
| 17 | 721.5779 | 745.5779 |
| 18 | 745.5888 | 769.5888 |
| 19 | 769.5997 | 797.5997 |
| 20 | 797.6124 | 827.6124 |
| 21 | 827.6261 | 861.6261 |
| 22 | 861.6415 | 901.6415 |
| 23 | 901.6597 | 957.6597 |
| 24 | 957.6852 | 1,041.69 |
| 25 | 1,041.72 | 1,249.72 |

